# Climate change and green sea turtle sex ratio – Preventing possible extinction

**DOI:** 10.1101/2020.04.08.032227

**Authors:** Jana Blechschmidt, Meike J. Wittmann, Chantal Blüml

**Author notes:** Correspondence: J.B., C.B.

## Abstract

Climate change poses a threat to species with temperature-dependent sex determination. A recent study on green sea turtles (*Chelonia mydas)* at the northern Great Barrier Reef (GBR) showed a highly female-skewed sex ratio with almost all juvenile turtles being female. This shortage of males might eventually cause population extinction, unless rapid evolutionary rescue, migration or conservation efforts ensure a sufficient number of males. We built a stochastic individual-based model inspired by *C. mydas*, but potentially transferrable to other species with TSD. Nest depth, level of shade, and pivotal temperature were evolvable traits. Additionally, we considered the effect of crossbreeding between the northern and southern GBR, nest-site philopatry, and conservation efforts. Among the evolvable traits, nest depth was the most likely to rescue the population in the face of climate change, but even here the more extreme climate-change scenario led to extinction. Surprisingly, nest-site philopatry elevated extinction rates. Conservation efforts to artificially increase nest depth promoted population survival and did not preclude trait evolution. Although extra information is needed to make reliable predictions for the fate of green sea turtles, our results illustrate how evolution can shape the fate of long lived, vulnerable species in the face of climate change.

**Graphical Abstract:** 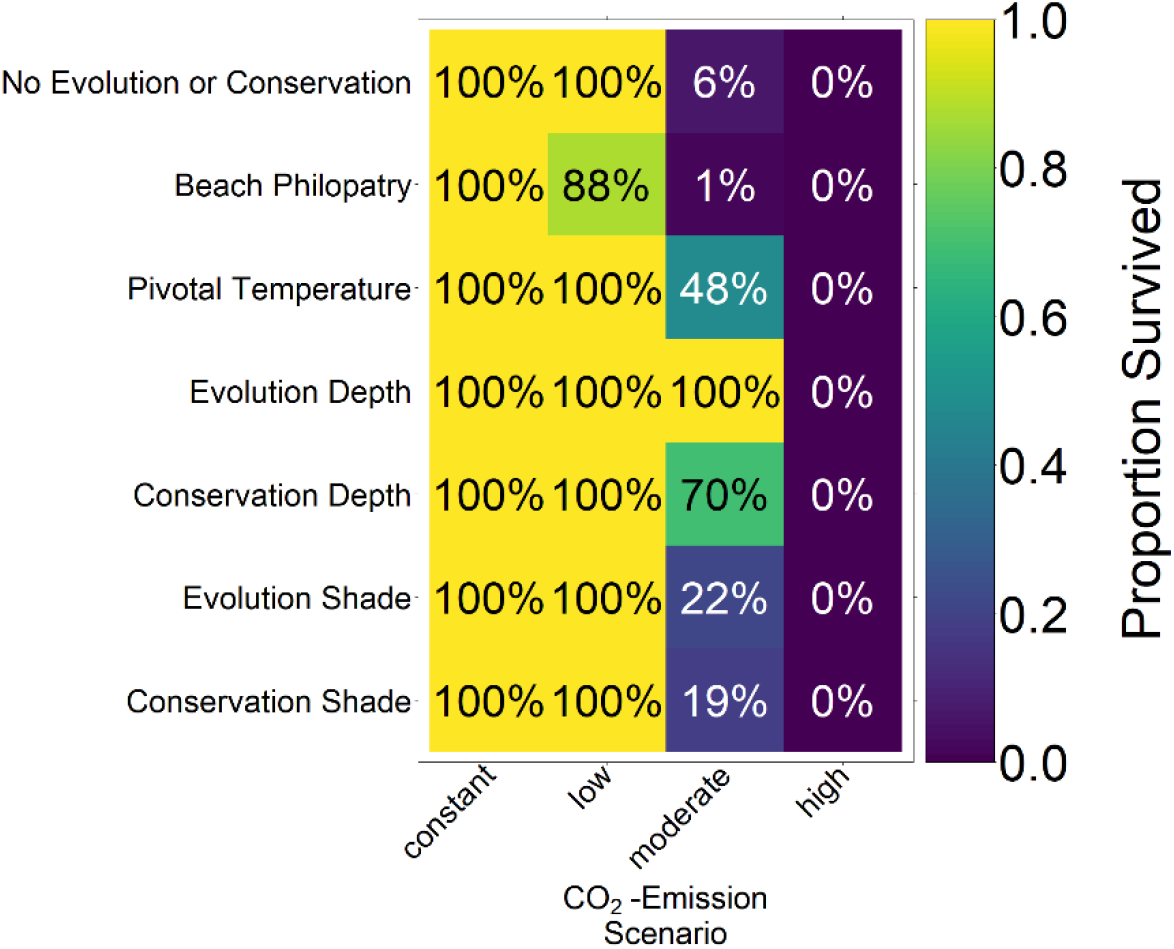

## 1. Introduction

Global warming poses a potential threat to biodiversity all over the world. A group of species at particular risk are long-lived reptile species with temperature-dependent sex-determination (TSD) [1,2]. In these species, increases in temperature can lead to biased sex ratios. Shortage of one sex is then expected to lead to mate-finding difficulties, failure to reproduce, and ultimately population decline [3]. This can be seen as a mate-finding Allee effect [4]. The persistence of these species will depend on whether or not they are able to adapt their sex-determination system rapidly enough with increasing temperatures to prevent extinction. In other words, such species may require evolutionary rescue [5] to persist.

The green sea turtle (*Chelonia mydas*) is an example for a long-lived species where survival of populations may depend on rapid adaptation to increasing temperatures [6]. Females express nest site philopatry, that is, they come back to the same beach where they were born and deposit four to five nests with about 100 eggs per nest, which are covered with sand [7,8]. The embryo’s sex is determined during the second third of incubation [9]. The warmer the egg, the higher the probability of a female hatchling. Furthermore, very high temperatures increase egg mortality [10]. The pivotal temperature is defined as the temperature at which 50% of the hatchlings are female. Pivotal temperatures vary across species and between populations [9,11,12]. Here we focus on the northern Great Barrier Reef (nGBR) population, which has a pivotal temperature of 29.3°C [13]. A slight change in temperature can alter the sex ratio of turtle hatchlings substantially. Just 1°C above the pivotal temperature, 80% of hatchlings are female [13]. Global mean temperatures, however, are expected to increase by 1.0°C to 3.7°C until 2100 [14]. Within the lifespan of an individual turtle, the temperature might have already increased by 0.8°C [14]. The oldest turtle fossil records date back to the Triassic (200 million years), indicating that turtles have survived several climate fluctuations in the past [15]. However, it is unclear whether the species will be able to keep up with the high pace of current climate change.

The nGBR population is one of largest breeding green sea turtle populations in the world. It has over 200,000 breeding females [16]. Currently this population has an approximate sex ratio of 80% female, with 99% of the non-adult turtles being female [16]. It has been suggested that a female-biased sex-ratio may even enhance population growth [17], since few males can fertilize many females. However, the low generation turnover rate may lead to a delayed impact of the lack of males. Eventually, the few remaining males may no longer be able to fertilize enough females to sustain the population, and the population might go extinct [18]. Unfortunately, there is currently not much information on how female fertilization probability depends on sex ratio or the number of males in the population. It is commonly assumed that these turtles mate in designated mating areas [19]. Moreover, while females can mate about every three years, and store sperm throughout a season, males are able mate every year [8,20,21].

A recent study suggests that juvenile recruitment in the nGBR population has decreased in the last few years [22]. Adaptation to higher temperatures could be necessary for this population to avoid extinction. Since evolutionary rescue is more likely in large populations [23], the nGBR population may be a good candidate. With increasing bias in the sex ratio, a trait controlling offspring sex ratio would be under strong selection pressure. Depending on the frequency of males and females within the population, selection would favour the less common one according to frequency-dependent selection [24]. For example, in a highly female-biased population, a trait leading to a production of more males would have an advantage. In green sea turtles, selection could act on a number of traits that influence egg incubation temperature or pivotal temperature and thus control hatchling sex ratio. We here consider four such traits:

1. Nest depth. Green sea turtle females bury their eggs anywhere from 30 to 90cm deep into the sand [7]. When comparing nests at different depths, deeper nests are on average cooler than more shallow nests [25]. The exact temperature depends on many factors, e.g. the beach vegetation, wind, and grain characteristics such as size, colour, and moisture [26–29]. Moreover, shallow nests experience stronger temperature fluctuations than deeper nests [25].
2. Shading of nest. On a typical nesting beach, there is substantial vegetation that provides shade throughout the day [7]. Depending on the hours of direct sunlight that the nest receives, temperatures vary [30]. The coolest nests are those directly underneath a tree or bush since they will be exposed to the sun for the shortest amount of time. Too much vegetation, however, can also be detrimental because roots may deter turtles from digging their nests [30]. The mean nest temperature at a medium level of shade (15%) is 1°C cooler than for nests located in the sun, and nests with a high level of shade (30%) were found to be 1.9°C cooler [30]. These temperature differences could potentially have an effect on hatchling sex ratio.
3. Pivotal temperature. The pivotal temperature varies between species with TSD and between populations of the same species [11]. Also within populations, heritable variation in the sex-determination threshold has been found in other turtle species [31]. The genetic basis of TSD is still a topic of research, and probably involves multiple loci. The best-studied locus is the cold-inducible RNA-binding protein (CIRBP) gene whose expression differs in embryonic gonads at different temperatures [32]. CIRBP has two alleles, one of which is thermosensitive while the other is not [32]. The expression pattern of CIRBP within developing gonads shows that it is able to mediate temperature effects on the bipotential gonads, effectively shifting the pivotal temperature [33]. Allele frequencies differ between snapping turtle populations and are correlated with pivotal temperatures [33]. This suggests potential for adaption. However, to our knowledge there is currently no data on CIRBP allele frequencies in *Chelonia mydas*.
4. Choice of nesting beach. Green sea turtles display maternal nest-site philopatry [7]. When a female reaches sexual maturity, she returns to her natal beach for oviposition. Different beaches can have different temperature conditions, depending on sand colour, grain size and vegetation. Additionally, the orientation of the beach on the island may also be important.

Since climate change is occurring at a very fast rate compared to historical temperature changes, evolutionary adaptations would have to occur rapidly to rescue green sea turtles at the nGBR [14]. Because of slow generation turnover, it has been doubted whether evolution can be fast enough [16]. Should the turtles not be able to adapt to the rising temperatures, artificially lowering nest temperatures may help to ensure population survival. Relocating nests deeper into the sand or providing them with shade should lead to an increase in male hatchlings. However, it is important that anthropogenic conservation efforts do not prevent evolution of the two traits in the long term.

Immigration of males from other populations may additionally promote the persistence of the nGBR population. The population at the southern Great Barrier Reef (sGBR) appears to be less susceptible to climate change than the northern one. It is located further away from the equator and hence does not experience equally high temperatures as the nGBR population. As a consequence, the sex ratio of the southern population is not as skewed as the northern one: it currently is at 67% female [16]. Males from the sGBR population could keep the nGBR population from experiencing a lack of males, even if the population consists exclusively of females [16]. It is known that some level of migration takes place between the two populations [21]. However, the extent of crossbreeding remains unknown.

We created a stochastic individual-based model that includes all the above-mentioned evolutionary traits as well as anthropogenic conservation efforts. Our goal is to analyse how nest depth, shading, beach philopatry, and pivotal temperature influence sex ratios and population size in the face of climate change. Additionally, we explored how conservation measures could be used to maintain the nGBR population. Lastly, we quantified the effect of crossbreeding between the nGBR and sGBR population.

## 2. Material and Methods

We built an individual-based stochastic model with overlapping generations. The model’s main purpose is to predict the survival probability and the sex ratio of the nGBR green sea turtle population for different climate, evolutionary, and conservation scenarios. In the first part of the study, we considered the fate of the population under constant temperatures between 25°C and 38°C. In the second part, we modelled four different temperature trajectories over the next centuries, based on the predictions of the Fifth Assessment Report of the Intergovernmental Panel on Climate Change (IPCC) [14,34]. The model comprises the timespan between 1800 and 2500.

The climate change model starts in 1800 to give the population time to equilibrate. For the first 110 years, the baseline nest temperature is kept constant at 29.3°C, which equals the pivotal temperature. This is based on the assumption that pre-industrial temperatures on average produced a 50-50 sex ratio. Starting in 1910 (year 111 in the simulation) when temperature data became available, real temperature recordings from the Australian Bureau of Meteorology [35] are fed into the model. Data for the region of Eastern Australia were used, since that includes both the sGBR and nGBR area. We used temperature data from December, the height of the breeding season [19]. We used 29.3°C as the baseline nest temperature, and then added the December mean temperature anomaly data. The temperature anomaly is here defined as the deviation from the average air temperature during 1961-1990, which is 25.8°C for this region. The average nest temperature is some degrees warmer, mainly due to metabolic heating of the nest [36,37]. We assumed that the anomaly in air temperature equals that in nest temperature and that the average baseline nest temperature during 1961-1990 continued to be 29.3°C.

From 2020 onwards, long-term predictions were used to determine the average temperature anomaly [14]. We used the four different IPCC Representative Concentration Pathways (RCP) scenarios RCP2.6, RCP4.5, RCP6, and RCP8.5. These are based on four different radiative forcing scenarios (the number representing W/m^2^ by 2100), each depending on the level of CO_2_ emissions (in the following referred to as constant, low, moderate, and high emission scenario).

The resulting average baseline nest temperature trajectories can be seen in Figure 1. In order to obtain realistic year-to-year temperature variability, we drew temperatures for all years without available weather data (before 1910 and from 2020 on) from a normal distribution. The means were the above-described temperatures for the respective years. For the standard deviation, we used the standard deviation of recorded temperatures from 1910 to 2019, which was 0.9532011. The standard deviation is indicated by shaded areas in Figure 1.

**Figure 1:**
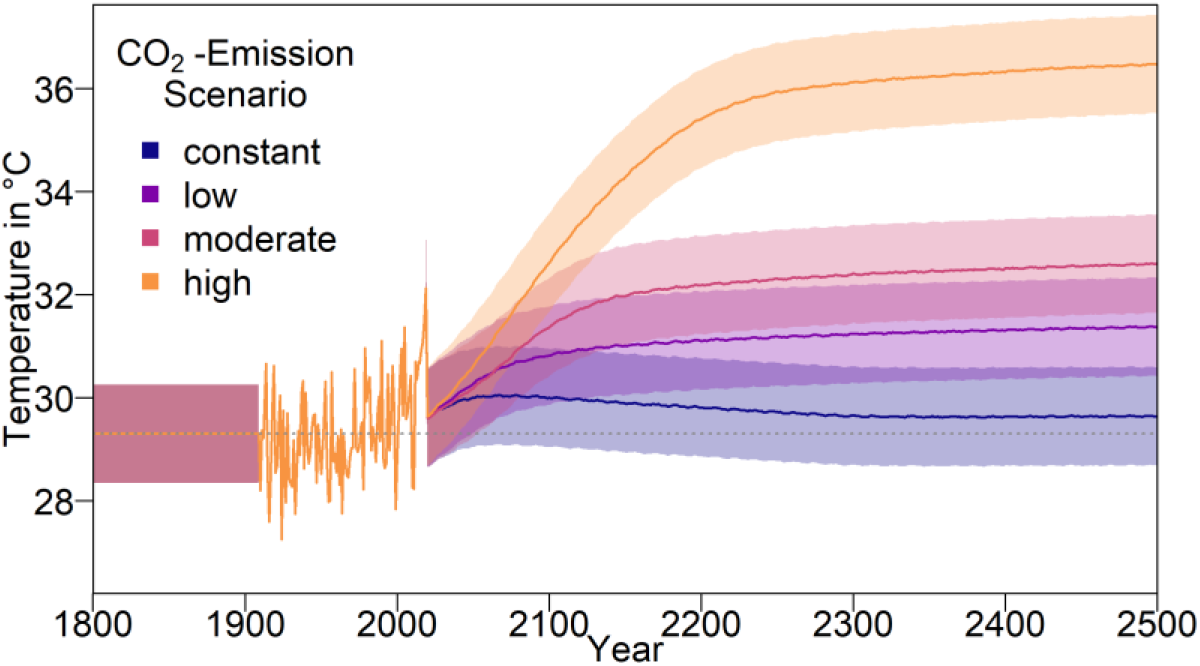
Temperature data used for the simulation. Before weather data became available (1800-1910), the average baseline nest temperature was assumed to be at 29.3°C (pivotal temperature for nGBR green sea turtle population). From 1910 to 2019, weather data was used. Values are derived from the average temperature anomaly (relative to the average temperature during 1961 to 1990) in Eastern Australia during December [35], which was added to the baseline temperature. Predicted temperatures starting in 2020 from are derived from the IPCC report [14,34]. Shaded areas indicate the standard deviation of weather data, which was used for drawing simulated temperatures during the implied intervals.

For each climate change scenario, we let the genes for nest depth, level of shade, pivotal temperature, or beach orientation evolve, as well as all combinations of those parameters with and without migration and conservation. We ran 100 replicates per setup.

The sex ratio of hatchlings as a function of nest temperature, *t*, is described by

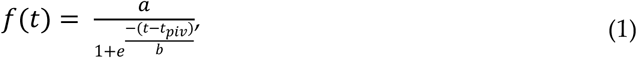

where *t*_*piv*_ = 29.3 is the pivotal temperature, *a* = 1, and *b* = 0.4424779. With these parameter values, the transitional range of temperatures (TRT), i.e. the difference between the temperature where 95% of hatchlings are female and the temperature where 5% of individuals are female, is 2.6°C. For details on parameter estimation see Appendix A.

Adult population size is variable but bounded by a carrying capacity *K*=200. We assume an initial population size *N_0_*=*K*. Note that for the sake of computational efficiently, our simulated populations are much smaller than the actual populations in nature. Supplementary simulations confirm that our results are robust to changes in carrying capacity and initial population size (see Appendix B, Figure A 1).

A female lays 100 eggs per breeding season if she found a mate and did not reproduce in the two prior years. Individual survival in the model is affected by several processes. Only 1.5% of hatchlings survive to the age of 40 [20], at which point they reach sexual maturity [16] and are included in our population size count. Note that in the wild, multiple nests with up to 100 eggs are common, but not all eggs hatch [13] and most of the eggs or hatchlings die before reaching maturity. What matters in our model is the number of offspring surviving to maturity. Since there is a lot of uncertainty in the literature regarding this number, we included the proportion of surviving offspring in a robustness analysis (Appendix B, Figure A 7). Additionally, the number of offspring is reduced according to carrying capacity by

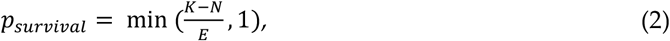

where *E* is the total number of juveniles that would reach maturity in a given year. Adult turtles have a yearly survival probability *p_adult_* = 0.9482 as estimated by Chaloupka [38]. In nature, egg survival furthermore decreases at extreme temperatures [10], which is however not included in the model.

Each female has a chance of meeting a male each year as long as she has not reproduced in the two preceding years [8,20,21]. There is currently little information on how males and females encounter each other. Here we assume that the probability for a female to encounter a male in a given year increases with the number of males in the population, *N_males_*, according to the equation

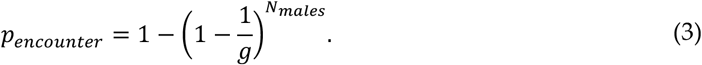

That is, the probability to not encounter a male decreases exponentially with increasing number of available males, with the rate of decrease determined by *g*. The parameter *g* can be interpreted as a number of mating areas if a male and female must randomly choose the same mating area in order to mate. *p_encounter_* can then be understood as 1 minus the probability that all *N_males_* males in the population go to a different mating area than the one chosen by the focal female. We chose *g*=100 as the default value, which would give an encounter probability of 0.63 at a population size of 200 with a 50-50 sex ratio. In Appendix B, Figure A 6, we explore the sensitivity of our results to the number of mating areas.

The genetics of the population are modelled as follows: Each individual has three diploid loci with two alleles each. There is one genetic locus each for nest depth, level of shade of the nest, and pivotal temperature, but only those relevant for the respective setup are taken into account to determine nest temperature and sex ratio. These traits are represented by real positive values between 0 and 1. Allele values for individuals in the initial population are independently drawn from a uniform distribution between 0 and 1. The mean of both of an individual’s alleles for a given trait is the phenotypically expressed trait for that individual. A phenotype of 0.3 for the nest depth trait for example would mean that the female will bury her eggs at a depth of 30cm.

Furthermore, each individual has a nesting beach, which depends on where it hatched, and therefore is a maternal effect.

The initial sex ratio of the starting population is 0.5 and individuals are evenly distributed over hatching beaches. Juveniles maturing within the first 40 years were generated in the same way as the rest of the starting population. This means each of these initial cohorts of juveniles has a 50/50 sex ratio and a third of them hatched on each beach. Ages of juveniles were uniformly distributed between 1 and 39. We generated *K*\*5 Juveniles per year for the model to draw offspring from within the first 39 years. From this pool, maturing animals were drawn during the first 39 years of the simulation. Each individual had a chance to be added to the adult population as described by equation 2. By running the simulation for a number of years prior to climate change, we ensured that the initial conditions did not influence the population’s response to climate change.

The genetic traits are inherited from both parents. For each trait, one of the mother’s alleles and one of the father’s alleles is randomly chosen for the offspring. When being passed on, alleles are susceptible to mutation. Mutation is implemented by drawing the new allele value from a Beta distribution. The two shape parameters, α and β, are calculated from the variance and mean of the distribution. The mean of the distribution, *μ*, is set to the original allele value. To our knowledge there is no empirical data on mutation processes in *C. mydas*, so we chose a standard deviation of 0.01, which leads to the variance *σ*^2^ = 0.0001. α and β then follow from

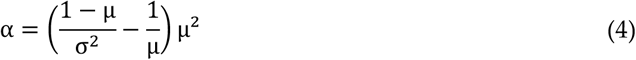

and

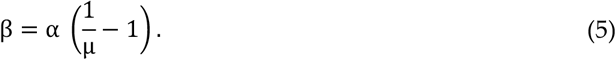

If the drawn trait value is below 0.001 or above 0.999, the value is set to these boundary values, as for values of 1 or 0, the beta-distribution would break down. For the beach orientation, each individual has a chance of ϱ=0.005 of changing their beach orientation trait value to one of the other two possible beach orientations. Results for an altered ϱ can be found in Appendix B Figure A 8.

Depending on the mother’s traits, the nest temperature will be warmer or colder than the temperature given by the environment. All effects of traits are added up and then added to the current year’s baseline nest temperature.

Results from measuring nest temperatures at differing depths [39] show a linear relationship with a decrease of 5.6 °C per m. We use this to determine the temperature difference Δ*t* compared to the environmental temperature:

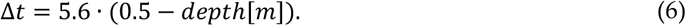

The depth of the nest is determined by an individual’s genetics as described above, allowing a maximum temperature difference of 5.6°C between the deepest and the shallowest nest. The default depth assumed in scenarios without depth evolution is 0.5m.

Secondly, a nest’s level of shade influences its temperature [30]. The temperature difference compared to the baseline temperature for the model can be described as

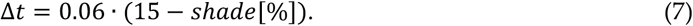

An individual’s preferred degree of shade for its nest will be determined by the two alleles in the same way as for the nest depth. Here, the genetic trait value, which is between 0 and 1, is multiplied by 30 to get the percentage of shade coverage. With 15% shade, nest temperature equals the year’s baseline temperature, 0% shade increases nest temperature by 0.9°C, and 30% shade reduces nest temperature by 0.9°C. Shading is limited to vary between 0-30% in the model, since that is the range covered by empirical data [30]. It might be possible to have higher levels of shade in nature, but the resulting nest temperature remains to be empirically investigated.

Furthermore, there might be a shift in pivotal temperature. This mechanism does not influence the nest temperature itself. Note that while the mother’s genes determine the depth and level of shade at which she buries the eggs, enzymes controlling offspring sex are produced within the egg. We therefore took the offspring’s genes into account to calculate its probability of developing into either sex. As illustrated in Figure 2, the same nest temperature will lead to different hatchling sex ratios, depending on the alleles. For a trait value of 0.5, the pivotal temperature is 29.3°C. In the model, the pivotal temperature can be shifted by at most 1°C in either direction for a trait value of 0 and 1. This is the average recorded difference in pivotal temperature between two populations of *Caretta caretta* in Australia [40]. We were unable to find any empirical data on variation in pivotal temperature within a population in *C. mydas*. For simulation runs without evolution of pivotal temperature, all individuals acted as if they had a trait value of 0.5. Throughout, we assume that the parameters *a* and *b* of equation (1) and thus the transitional range of temperatures are constant.

**Figure 2:**
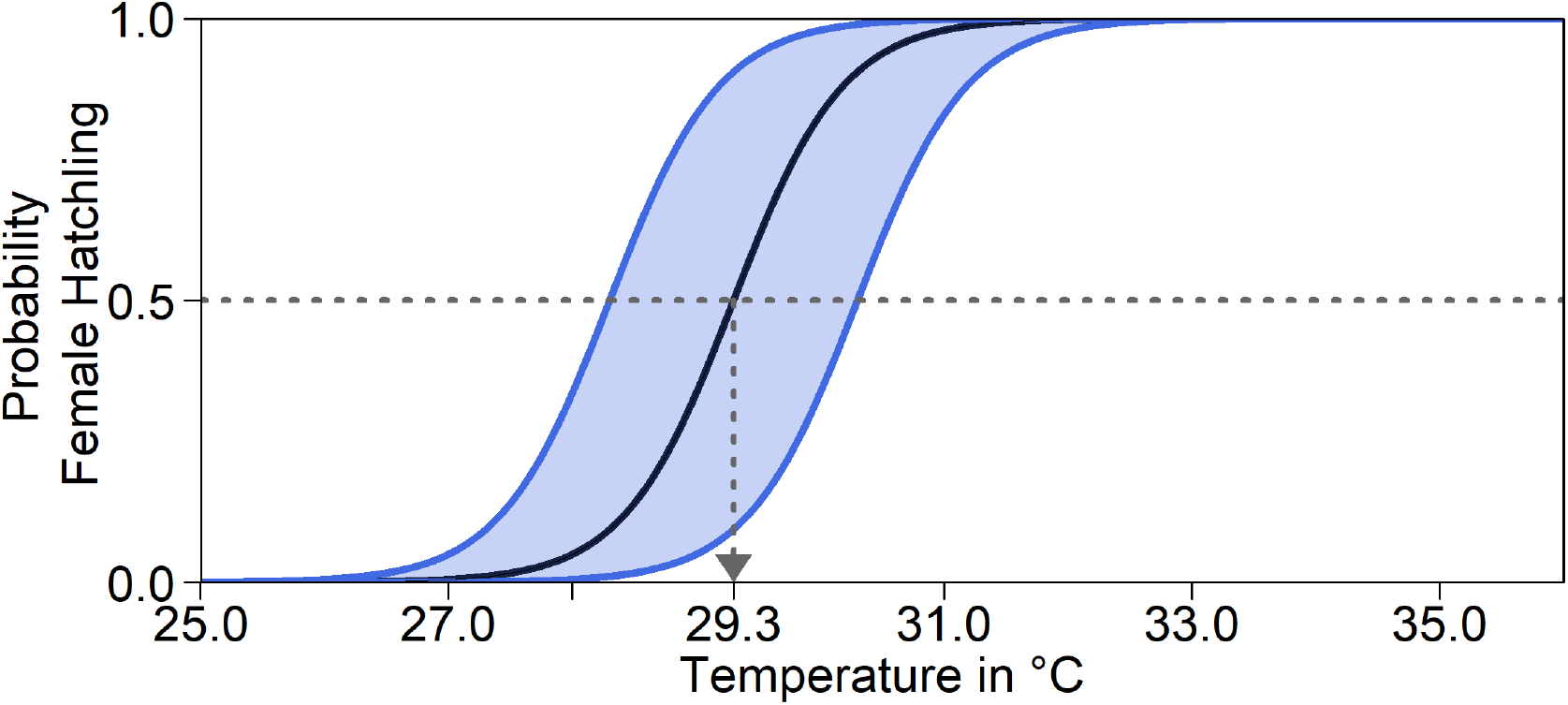
Probability of a female hatchling depending on temperature. The black line shows the result of equation (1) [13]. The pivotal temperature 29.3°C is indicated by the grey arrow. The blue lines are shifted by 1°C to the left or right, the maximum possible shift in out model. When the curve is shifted to the right, the pivotal temperature is higher, i.e. there are more male hatchlings at higher temperatures. When shifted to the left, pivotal temperature is reduced, and more females hatch at colder temperatures. The shaded area indicates the range used in the model.

Lastly, nest temperature is also influenced by beach orientation. Beach temperatures differ and are based on Booth & Freeman who measured nest temperatures on three beaches on Heron Island, which is located in the sGBR region [41]. In our model, we assume that temperature of nests on the eastern beach represents the average baseline nest temperature for that year. We assigned each individual one of the nesting beach orientations based on their beach of birth. Females later return there, and the corresponding temperature is then applied to their nests, altering the temperature by +0.7°C (north), 0°C (east), or −0.9°C (south), respectively [41].

As level of shade and nest depth seem easiest to manipulate in the field, we also included a setup where we tested the influence of conservation efforts on them. In the corresponding simulations, nest depth is changed to 0.9m, independently of the genetics; and shade level is increased to 30%. We decided to manipulate every second nest (*Ψ*=2) every ten years (*ϒ*=10) as default condition, starting in 2020. These parameters were also altered, and results can be found in Appendix B, Figure A 9.

The probability of meeting a male from the sGBR is constant over time. Due to a lack of information about the extent of crossbreeding between populations, we simulated a range of different meeting probabilities. If a female encounters males from both populations, the nGBR male will fertilize all eggs. For the model, we created a sGBR population under the same conditions as the nGBR population, but always with a constant temperature of 29.61336°C that would result in their current sex ratio of 0.67 [16] in the absence of evolution or conservation measures. Since the sGBR’s setup matched the one of the nGBR population they migrated to, they were potentially able to reach a 50/50 sex ratio again by evolving accordingly. This population was simulated for 1000 years to allow it to equilibrate. We collected all adult males from 100 simulations per setup in order to have a large pool of males available to migrate to the nGBR populations. Not all populations survived or had an equal proportion of males, so the total number of males to choose from differs between setups (Appendix B, Figure A 10).

Note that we tested the effectiveness of each evolutionary mechanism individually at first, without the influence of conservation and/or migration.

## 3. Results

### 3.1 Constant temperatures

Keeping the temperatures constant over the timespan of the simulation and allowing no evolution leads to at least 95% females for temperatures above 30.6°C. Figure 3 shows average population size (a) and average sex ratio (b) over time. For both high and low temperatures, population size decreases over time while it stays roughly constant at temperatures around the pivotal temperature. Sex ratios are strongly dependent on temperatures.

**Figure 3:**
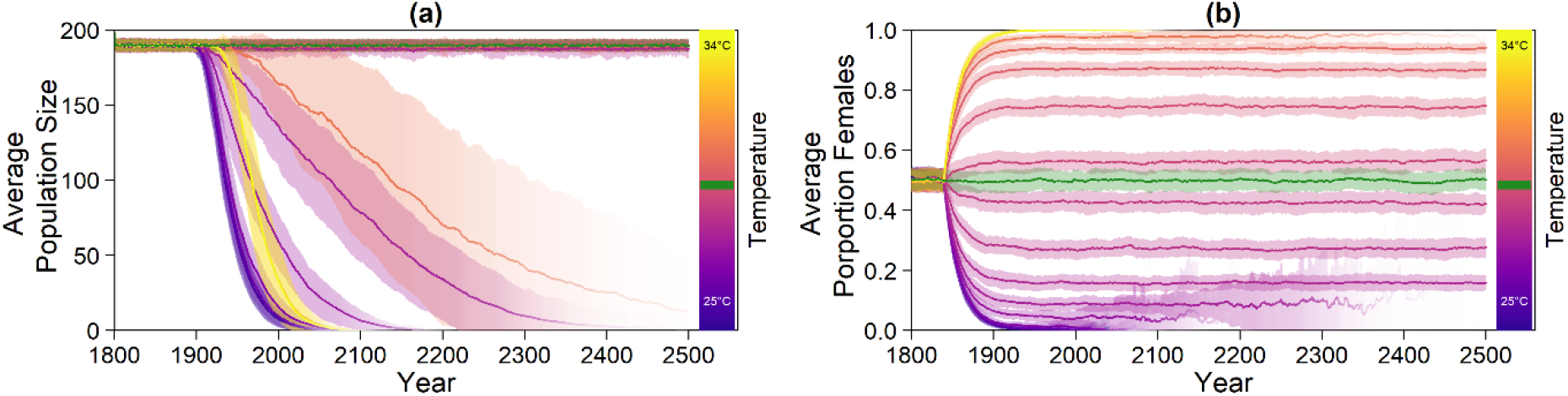
Model results without evolution of any trait and at various constant temperatures (different colours). (a) Average population size; (b) Average proportion of females. Colour intensity indicates the number of populations that have not gone extinct. While extinct populations are included as zeros in population size averages, they are excluded from the calculation of the average proportion of females. The green line indicates replicates run at the pivotal temperature of 29.3°C. Shaded areas show the standard deviation among replicates.

We simulated populations with all four possible evolutionary mechanisms: nest depth, pivotal temperature, level of shade, and beach orientation, and their combinations (Figure 4). All mechanisms except beach orientation increase average population sizes at the end of the simulation for both colder and warmer temperatures. The most effective setup for survival at high temperatures is the combination of all evolutionary mechanisms. Among the single mechanisms, nest depth promotes population survival the most. Beach orientation is the least effective evolutionary mechanism for adapting to warmer temperatures.

**Figure 4:**
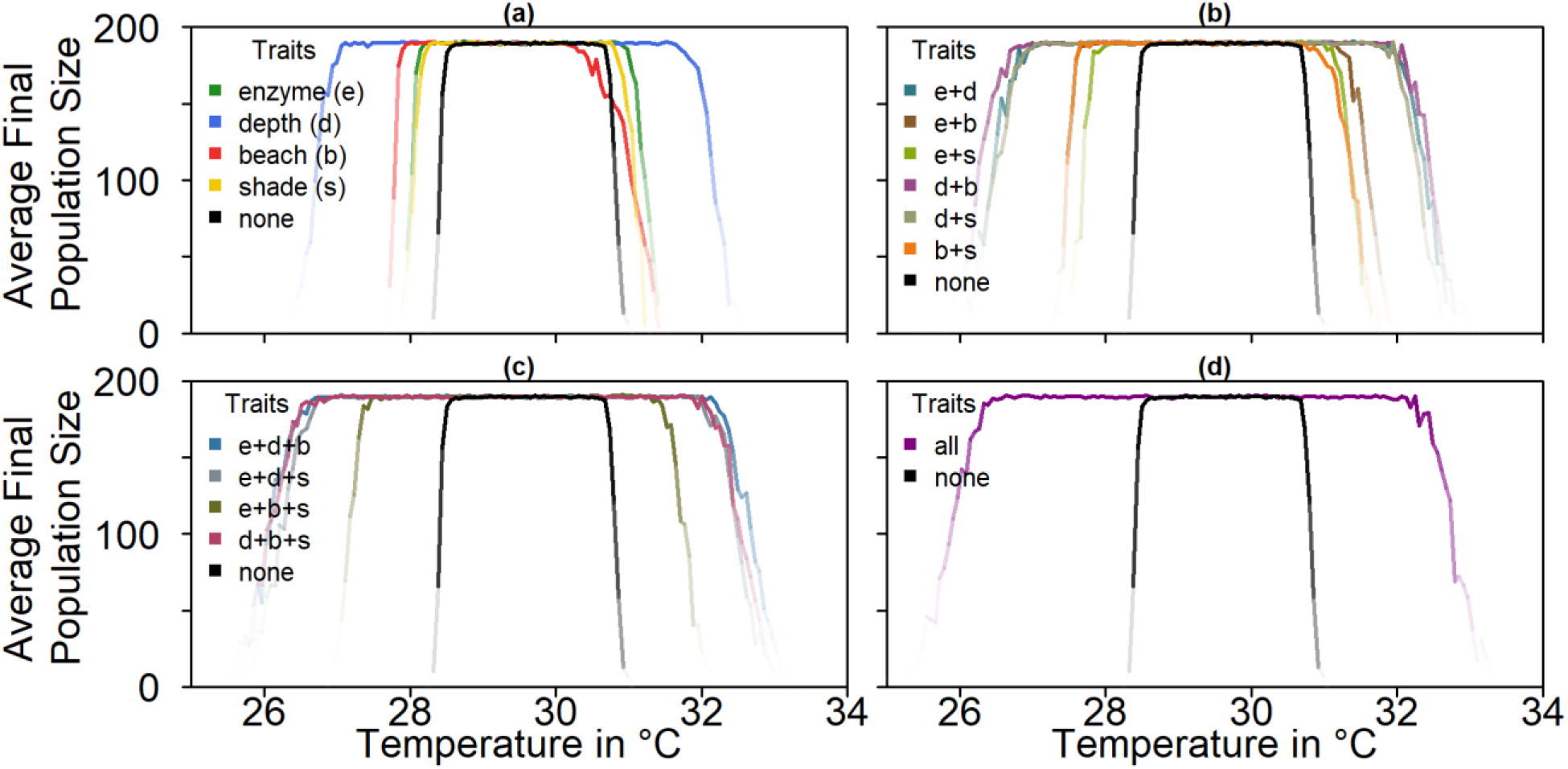
Average population size over 100 replicates at the end of the simulation (year 2500) depending on temperature, comparing all mechanisms and any combination of them. The black lines represent average population sizes without any evolution; coloured lines represent different evolutionary mechanisms. Colour intensity indicates the number of populations that have not gone extinct: (a) One trait; (b) Combination of two traits; (c) Combination of three traits; (d) Combination of all four traits.

### 3.2 Climate Change

For the climate-change simulation, there are four different scenarios (see Figure 1). Predictions for average population sizes and sex ratios with no adaptation to climate change are shown in Figure 5.

**Figure 5:**
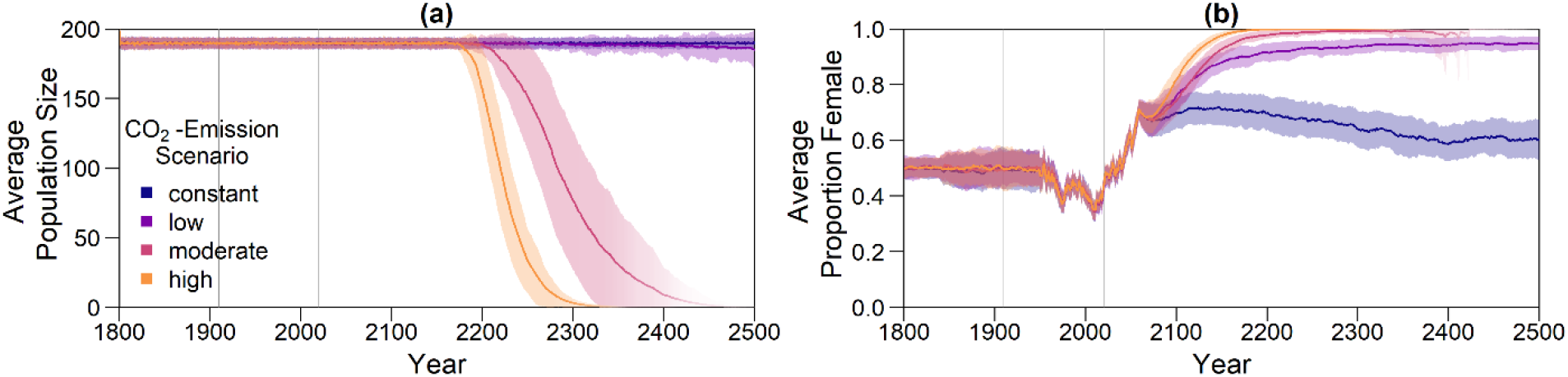
Results for the climate change simulation without evolvable traits. Intensity of the line represents proportion of surviving populations. Shaded areas indicate the standard deviation between replicates. Grey lines mark the points in time from which on weather data (1910) and temperature predictions (2020) were used. Note that the effect of the change in temperature is visible in the sex ratio with a 40-year delay, as only adult individuals are taken into account for averaging. (a) Average adult population size over time; (b) Proportion of females among adults over time. Each line in (b) represents the average of all non-extinct replicates at the respective time point.

Without evolution, population size drops dramatically within the next centuries for the moderate and high emission scenario, with the fastest decline for the high emission scenario (Figure 5a). For the moderate and high scenario, almost all populations go extinct (Figure 6). With a constant and low level of CO_2_ emissions, the population size stays stable close to carrying capacity. Figure 5b illustrates the sex ratio of the population over the years. Starting in the year 1910, when weather recordings were incorporated, sex ratio begins fluctuating and eventually becomes more female skewed when juveniles born during 1990 to 2018 reach maturity 40 years later. While for the low emission scenario, the average sex ratio is 0.95, the moderate and high scenarios reach a proportion of females of around 1. In the constant emission scenario, a 0.6 female ratio is reached and maintained.

**Figure 6:**
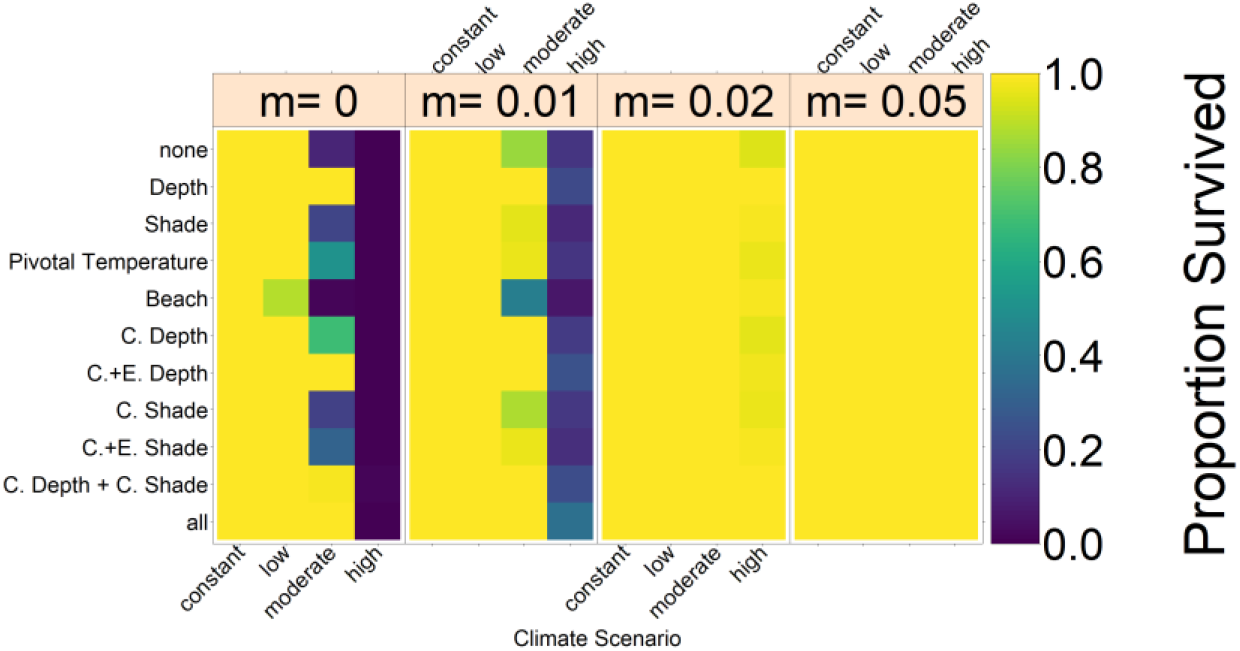
Overview of the proportion of populations in the various setups that did not go extinct by 2500. The left block represents the results without migration and the blocks to the right the results for different migration rates, *m* (indicated in top box, see section 3.2.3). Colours indicate the proportion of populations that still persist in 2500. Each row depicts one evolutionary mechanism (C=conservation, E=evolution). Each column shows the result for one climate scenario. Sections show results for different migration rates between 0 and 0.05. Proportion of surviving populations are indicated by colours.

Note that the actual sex ratio of adult nGBR turtles in 2018 is 0.86 female, while it is closer to 0.38 in 2018 in the model. This might be due to the fact that we selected 29.3°C as the average baseline nest temperature from 1961-1990, when it might have been already warmer than that for this time period. However, the discrepancy may also be due to the fact that we define “adult turtles” as turtles that are 40 years and older, while Jensen et al. group turtles based on size, so that they count them as adults around the age of 25 [16]. Adult turtles in 2018 in our model therefore had their sex determined before 1978, when the temperature anomaly from weather data were more often negative (leading to more males) than positive [35]. Since there are more males among older individuals, this could skew our sex ratio towards males compared to their study.

#### 3.2.1 Evolution of each trait

Figure 7 shows the results for the evolution of nest depth, level of shade and pivotal temperature. Evolution of nest depth is the most effective mechanism, keeping the population stable close to carrying capacity for the constant, low, and moderate emission scenario. For evolution of level of shade and pivotal temperature, the constant and low scenarios are survived as well, while a large proportion of populations go extinct for the moderate one (Figure 6). The sex ratios fluctuate according to the recorded temperatures in a similar way for all climate scenarios before changing according to the climate scenario and the respective evolutionary mechanism. After an initial rise in sex ratio starting in 2020, the increase in females can be reversed by evolution of traits (lower row). Average nest depth, level of shade and pivotal temperature all increase for the three warmer emission scenarios. For the warmest scenarios, a sex ratio of 1 is reached before evolution of the respective trait can counteract the feminization.

**Figure 7:**
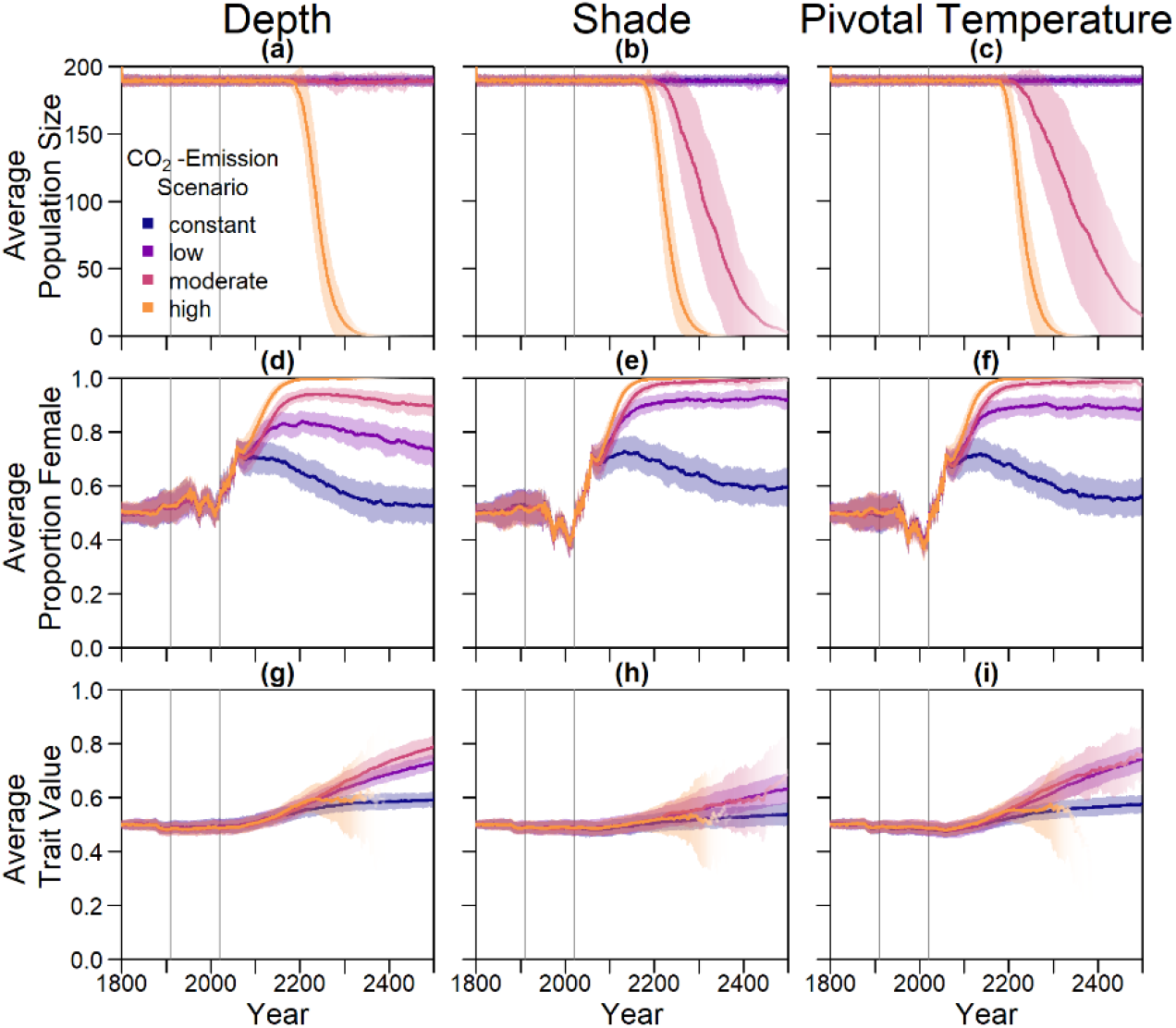
Results for evolution of nest depth, level of shade and pivotal temperature. Intensity of the line represents proportion of surviving populations. Shaded areas indicate the standard deviation between replicates. Grey lines mark the points in time from which on weather data (1910) and temperature predictions (2020) were used. Note that many effects are visible with a 40-year delay, as only adult individuals are taken into account for averaging. (a-c) Average population size over time; (d-f) Average proportion of females among adults over time; (g-i) Average genetic traits for nest depth, level of shade eggs are laid in, and pivotal temperature shift over time.

**Figure 8.**
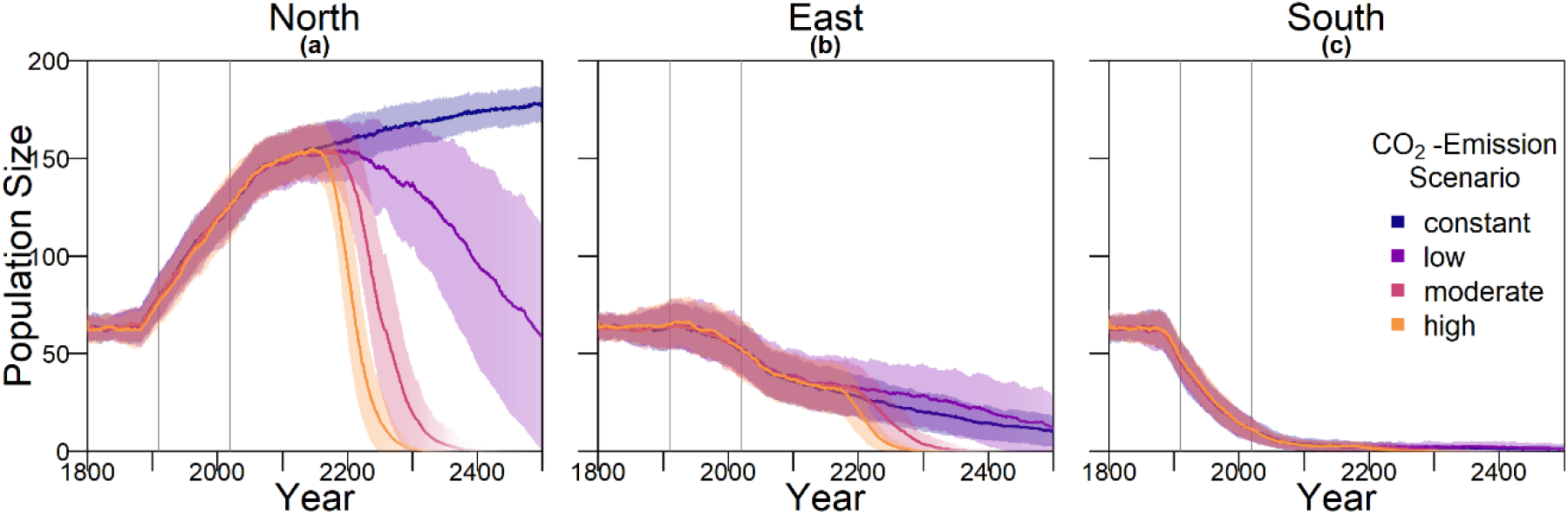
Population size on each of the nesting beaches. Intensity of the line represents proportion of surviving populations. Shaded areas indicate the standard deviation between replicates. Grey lines mark the points in time from which on weather data (1910) and temperature predictions (2020) were used. Note that the effect of a change in temperature is only visible in the sex ratio with a 40-year delay, as only adult individuals are taken into account for averaging: (a) Average population size on the northern beach, which is 0.7°C warmer than the eastern beach; (b) Average population size on the eastern beach, which is assumed to correspond to the respective baseline nest temperature; (c) Average population size on the southern beach, which is 0.9°C colder than the eastern beach.

The evolution of the beach orientation leads to extinction in all replicates for the two warmer climate scenarios (Figure 6). Population size drops on the eastern and southern beaches, while it increases on the warmest, northern beach, before the population goes extinct. In the low emission scenario, 88% of populations persist until 2500, but the average population size strongly declines to an average of around 50 at the end of the simulation. In contrast to the other evolutionary scenarios where long-term persistence appears possible in the low emission scenario, all populations with the beach evolutionary mechanism would likely go extinct after 2500.

As under constant temperature, we also explored combinations of evolutionary mechanisms under climate change. An overview of population persistence in these can be found in Appendix B Figure A 6. Generally, the more evolutionary mechanisms are combined, the higher the proportion of survival. Again, there is one exception: The evolution of the beach orientation leads to a slightly higher extinction rate among populations.

In Appendix B, we explore the robustness of our results to changes in various parameters. The effect of different numbers of mating areas is shown in Appendix B, Figure A 6. Population survival prospects decrease with an increasing number of mating areas, regardless of the evolutionary mechanism at hand. For fewer mating areas, population survival depends greatly on the chosen evolutionary mechanism. Additionally, we explored the sensitivity of our results to the following parameters: age at maturity (Figure A 2), survival rate (Figure A 3), mutation rate (Figure A 4), and pre-industrial nest temperature (Figure A 5). All parameters were varied by ±5%. The model was most sensitive to changes in survival rates. When increasing survival rate of both juveniles and adults by 5%, proportion of populations surviving to 2500 reaches 100% for all possible scenarios and mechanisms, including the high emission one. This is because the yearly probability for an individual to die is then only 0.439%, making the average adult lifespan 227.79 years compared to 19.3 years (age of 59.3) in our default setup. Consequently, only 3 generations pass over the course of the simulation. Lowering survival rate by 5% did not have the same extreme effect, but population survival did decrease for all scenarios. For all other parameters, varying them did not lead to any unexpected or large differences in population survival. Moreover, we altered juvenile survival, i.e. the survival probability from egg to mature adult (Appendix B Figure A 7). The model is most sensitive to changes in juvenile survival between 0.001 and 0.02. Increasing survival leads to an increase in overall population survival probability for most emission scenarios. No juvenile survival rate that we tested enabled the population to survive the high emission scenario.

#### 3.2.2 Conservation efforts

Figure 9 shows the results of conservation efforts, without and in combination with trait evolution. Conservation efforts are carried out every 10 years and are applied to every second nest (for other parameter combinations see Appendix B, Figure A 9). Nests are put at a depth of 0.9m and at a 30% level of shade, respectively.

**Figure 9:**
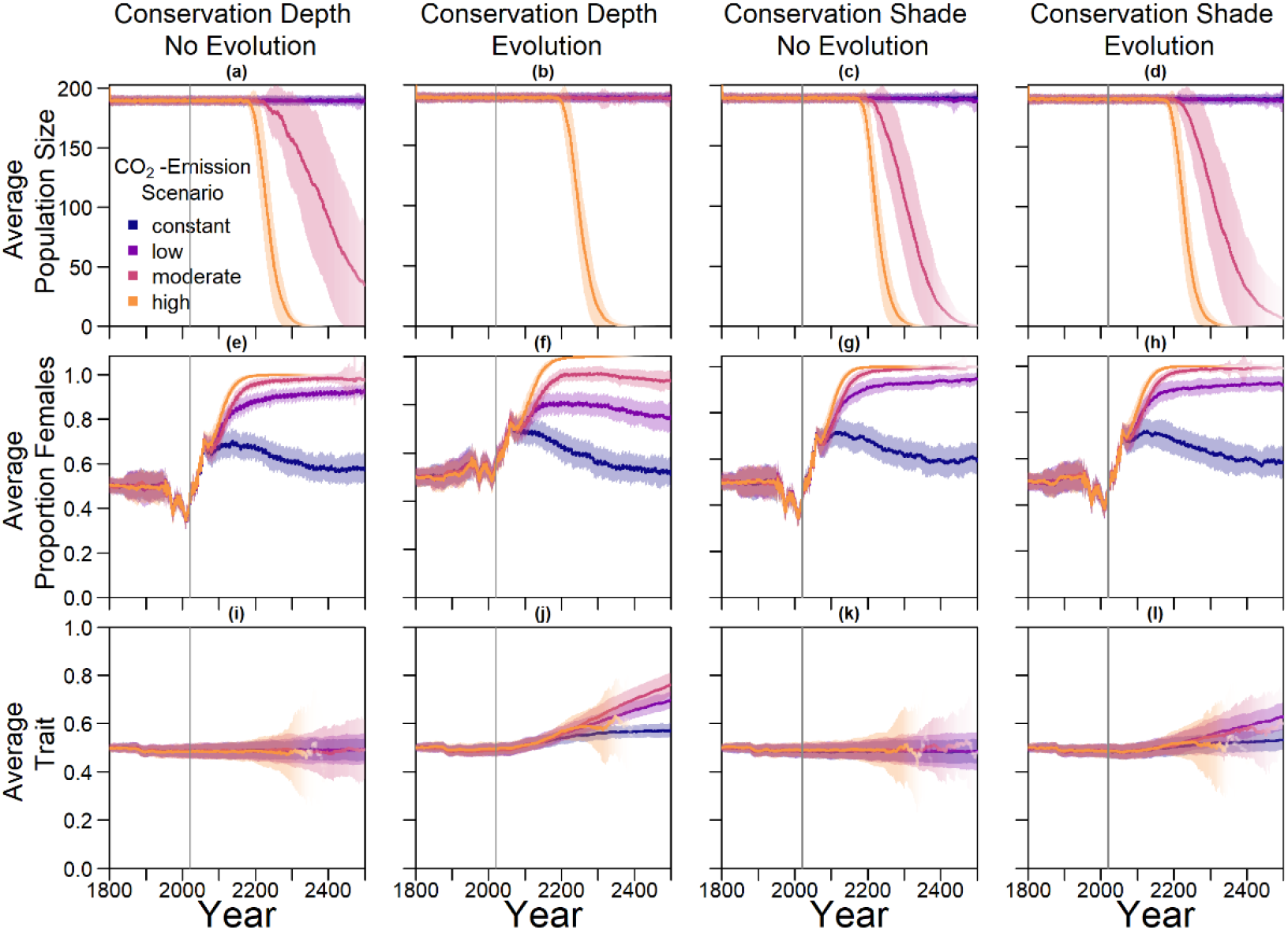
Results for conservation efforts combined with evolution of level of shade and nest depth. Gray lines indicate the start of conservation measures in 2020. (a) Average population size over time where every 2^nd^ nest is moved to 90cm every 10 years, but nest depth is not an evolvable trait; (b) Average population size over time where every 2^nd^ nest is moved to 90cm every 10 years, and nest depth is an evolvable trait; (c) Average population size over time where every 2^nd^ nest is moved to 30% shade every 10 years, but preferred level of shade is not an evolvable trait; (d) Average population size over time where every 2^nd^ nest is moved to 30% shade every 10 years, but preferred level of shade is an evolvable trait; (e-f) Corresponding average proportions of females among adults over time; (i-l) Corresponding average trait values over time.

On the left-hand side of Figure 9, we present the population size and sex ratio over time for conservation by manipulating nest depth with and without evolution. For the moderate and high emission scenario, population size declines over time without evolution, while for the constant and low scenario it stays at capacity. The higher the emission scenario, the quicker the decline. For the high emission scenario, population size drops to zero before the year 2400. When combining conservation and evolution, average population size stays close to carrying capacity for the constant and low scenario. Nest depth conservation alone increases the proportion of populations that survive until 2500 from 6% to 70% in the moderate emission scenario (Figure 6). When this trait additionally is evolvable, population survival increases to 100%. This pattern repeats less pronounced for shade conservation in the moderate emission scenario, where 94% of populations went extinct without evolution or conservation, but with conservation efforts alone the proportion of surviving populations reaches 19% and combined with evolution 27%. Only for the high emission scenario, even the combination of conservation and evolution does not lead to any populations surviving to 2500.

Accordingly, sex ratio is less female skewed when combining conservation and evolution. The average trait value stays at 0.5 on average for nest depth and level of shade when there is no selection on traits, as expected. When combining conservation and evolution, the trait evolves towards similar levels as without conservation (Figure 7).

Results for different rates of conservation measures are shown in Appendix B, Figure A 9. The more nests the measures are applied to, the higher the proportion of surviving populations. Additionally, the more often an effort is carried out, the more effective it is. The most effective combination that we tested is carrying the measures out every two years and applying them to every second nest. We also simulated combinations of both conservation measures at the same time. Generally, when combining the evolution and conservation of nest depth and level of shade, the population survives all climate scenarios except the high one, no matter the degree of conservation. Even when only applying measures every 20 years to every 20^th^ nest, these three climate scenarios lead to a survival probability of 1. The high emission scenario, however, always has a survival probability of 0, even when measures are applied every second year to every second nest (see Appendix B, Figure A 9).

#### 3.2.3 Migration

For this part of the model, we created a southern GBR population that was used as a mating pool for the nGBR females. When creating these mating pools, not every scenario led to the same number of males in the southern GBR population (Appendix B, Figure A 10). The main cause of reduction in population size was the choice of nesting beach, just as for the northern GBR population. However, for every scenario there was always a sufficient number of males to serve as immigrants to the nGBR population.

We tested different levels of migration between the northern and southern Great Barrier Reef populations. The migration parameter rate *m* is defined as the probability for a northern female to meet a southern male in a given year. A higher migration rate leads to an increased chance of population persistence for all evolutionary mechanisms and climate scenarios (Figure 6). Even small migration rates drastically improve the proportion of populations that survive until 2500 under all climate scenarios and for all schemes. For a migration rate of 0.05 or higher, 100% of populations survived in all setups.

## 4. Discussion

The predictions for the four future climate scenarios without any evolution show daunting results. The two higher CO_2_ emission scenarios lead to extinction of the nGBR green sea turtle population. Currently, surface temperature has already increased by 0.8°C compared to pre-industrial temperatures [14]. According to our model predictions, a further increase by 0.5°C, elevating baseline nest temperatures to 30.6°C, poses a threat to population survival (see Figure 4). However, our results suggest that evolution or conservation efforts might enable green sea turtles to recover from their current strongly skewed sex ratio, at least under the less extreme CO_2_ emission scenarios.

Based on our assumptions, evolution of nest depth appears to be the most promising route to evolutionary rescue. A major reason might be that it allows the largest reduction of nest temperature in our model (up to 2.8°C), compared to the other traits that were assessed. With nest-depth evolution, simulated populations survive the constant, low, and moderate CO_2_ emission scenarios. Evolution of the trait will counteract the shift in sex ratio towards females. For the high emission scenario, the temperature rises too quickly for evolution to keep pace. The sex ratio then reaches 100% female and the population goes extinct before a shift in nest depth might produce more males. Compared to nest depth, evolution of pivotal temperature and nest shading had similar yet weaker beneficial effects.

Contrary to our hypothesis that the option to deposit eggs on colder beaches would enhance population persistence, the maternal effect of nest site philopatry made the turtles go extinct faster than without any evolving traits. Northern beaches are closer to the equator and therefore the warmest, leading to more females being born there. Females from northern beaches will mate with males from southern beaches, but offspring will lay their eggs on the northern beaches as well. It is a vicious circle: More turtles on northern beaches leads to more females, leads to more eggs on northern beaches, leads to more females, reducing male density even faster. Such a runaway process was already suggested by Bull [42]. Via a similar effect, maternally inherited nest-site choice hindered adaptation to climate change in a mathematical model for painted turtle (*Chrysemys picta*) populations [43]. Indeed, it has been proposed that maternally inherited nest-site choice can be one of the factors driving the evolution of environmental sex determination [44].

While the level of philopatry in green sea turtles is generally considered extremely high (93-97% for the Sarawak population), some individuals do not return to their native beach [7]. For the model, increasing “mutations” of the nesting beach trait increased the proportion of surviving populations (see Appendix B, Figure A 8). With a higher level of error, the possibility for having a few nests in colder temperatures and thus more males increases.

On a larger scale, there may be a similar effect between northern and southern GBR populations. Currently, there is no consensus on how frequent mating between the two populations is, but data suggest it might be quite common [21]. Since the southern population has a higher male ratio than the northern one, this could cause an increase in northern population size. A meeting frequency between 0.05 would, according to our model, largely increase the chance of population survival of the nGBR population in all emission scenarios. In loggerhead sea turtles in North America, there seems to be a similar relationship between northern and southern populations. While the southern populations are highly female skewed there, the northern ones could potentially provide them with males [45].

Our model results also show some interesting possibilities for conservation efforts. Putting the nests deeper into the ground by hand every few years might shift the sex ratio enough to ensure a ratio of males that is high enough to sustain the population. Shading nests may, however, be easier to implement. Unfortunately, in the model, it was not as effective as altering the nest depth. In the data used for nest temperatures under different levels of shade, the maximum amount of shade possible was at 30% [30]. Possibly more shade could be provided to further lower the temperature. However, implementing sun protection may prevent rainfall from reaching the nest. This will then make the nest warmer than its surroundings instead of colder [30], so any conservation must be implemented carefully and underlie constant supervision to ensure temperatures are actually lowered.

The model suggests that artificially changing nest depth or shade level does not interfere much with the natural evolution of these traits, e. g. nest depth still increases with increasing temperatures. However, the average nest depth trait value at the end of the simulation decreases by 2.8% when in combination with conservation compared to evolution only. This suggests that anthropogenic conservation efforts reduce selection pressure on the trait, but only to a very small degree, which may also be due to stochastic effects. For level of shade, the trait even increases by 0.5% relative to the scenario without evolution, which is likely due to noise.

Although the evolution of nest depth shows promising results for the moderate CO_2_ emission scenario, there is one big caveat: we do not know whether nest depth is a hereditary trait at all, since there is currently no data available on variation in nest depth between individual turtles in a population. One study suggests that nest depth may be correlated with limb size, so that larger turtles generally dig deeper nests [46]. Thus, if body size is a heritable trait, then nest depth might be as well. However, even if nest depth is a hereditary trait, more factors play an important role in determining the temperature of the nest besides the depth. The grain size of the sand on a nesting beach as well as its colour, the moisture of the ground, and levels of rainfall and shade all influence the temperature of nests. For example, in loggerhead sea turtles, high humidity seems to lead to higher male frequencies even at high temperatures [47]. Moreover, the temperature of the ground depends on the climate in the months before the mating season. Towards the end of the mating season, the ground is heated up by the summer [41]. As temperatures drop, the ground stays warmer, so in this case, the deeper the nest is in the ground, the warmer it is. In addition to these factors, nest depth might be limited by the ability of hatchlings to reach the surface and the durability of eggs themselves.

For pivotal temperatures, data for other turtle species suggest that there is substantial heritable variation within populations [31]. One of the genes underlying pivotal temperature is the cold-inducible RNA-binding protein (CIRBP) [33]. Change of CIRBP allele frequencies might be a likely route for evolutionary rescue of the nGBR green sea turtle population. The way the algorithm is set up in the model allows for the population to shift its pivotal temperature by up to one degree Celsius. It has been found before that CIRBP frequencies vary between populations of the same species, so it seems to be susceptible to evolution [33].

For the other evolving traits in our model, not much is known about the genetic basis and heritability. To improve the accuracy of the model, it is necessary to establish more data on these traits. Field studies on green sea turtles unfortunately take a long time to produce data [13].

In addition to the traits considered here, green sea turtles could potentially shift their breeding season in response to climate warming. In some bird species, a warmer climate has led to an earlier onset of breeding seasons [48]. For other turtle species, variation in nesting phenology has been documented across space and time [2,49,50]. During the end of spring, the ground is not as heated as during summer [41]. Thus, a shift in breeding season might mitigate the effects of climate change on the sex ratio of the turtles.

In both painted turtles, *Chrysemys picta*, and loggerhead sea turtles, *Caretta caretta*, females showed plasticity in the date of first nesting depending on year-to-year climatic variability [51,52]. Warmer winters lead to an earlier onset of nesting. A possible mechanistic explanation for a change in nesting date might be that a warmer winter allows *Chrysemys picta* to emerge earlier from hibernation, or provide them with more basking opportunities and thus influence the rate of egg development [51]. Another study on *Chrysemys picta* suggests that the level of shade provided for a nest might be a behavioural plastic trait [46].

All of these studies suggest that behavioural plasticity might be a possible way for species to alleviate the effects of rising temperatures. However, green sea turtle nesting dates do not seem to respond as much to year-to-year variation in sea surface temperatures [52]. This might be because of their different diet or because they only seldomly display hibernating [53] or basking behaviour [54]. Furthermore, the fact that nGBR green sea turtle sex ratios have already become heavily female-biased [16] suggests that green sea turtles have limited phenotypic plasticity for nest-site choice or nesting date in response to temperature.

Our results are based on historical data and predictions for average December temperatures and we do take into account interannual variation in mean December temperatures. Future modelling efforts could also include finer-scale temporal variation and small-scale spatial variation in temperatures. If at least some nests experience cold enough temperatures during the critical developmental time window, the resulting males could have a strong effect on the population’s persistence time. On the other hand, temperature fluctuations have also been suggested to have a feminizing effect, at least in painted turtles and red-eared slider turtles [31,55].

Another aspect that might potentially influence population survival but is not included in the model are the effects of inbreeding and small population size. When males become scarce and the same few males mate with most females, offspring are going to be more related than usual. This could lead to inbreeding depression and loss of genetic variation and evolutionary potential. Additionally, there is currently no data available as to how many females a male is able to mate with in nature and how the mate finding process works and depends on the density of males and females. Filling these gaps in knowledge could greatly improve the precision of model predictions in the future.

Several previous studies have attempted to make predictions for the fate of green sea turtles and other species with temperature-dependent sex determination under climate change. A study by Fuentes and Porter [56] compares two different modelling approaches to predicting soil temperatures in green sea turtle nesting grounds at the nGBR: a correlative model and a more complex and mechanistic microclimate model that takes into account climate maximum and minimum data (e.g. air temperature, wind speed, humidity, percentage of cloud cover) for arbitrary time intervals, e.g. monthly, weekly or daily, and physical properties of the soil (e.g. thermal conductivity, density, specific heat and substrate reflectivity). In the correlative model, they find that using both air and sea surface temperatures to predict soil temperature leads to a more accurate prediction of soil temperature than using either air or sea surface temperature on their own [29]. They find that microclimate and correlative modelling approaches lead to different soil temperatures and therefore different sex ratio predictions for the nGBR population, but agree in that a complete feminization is inevitable should temperatures keep rising as predicted by most emission scenarios by the IPCC [14]. However, they assume that the sex-determination system remains constant over time and does not evolve. A productive direction for future work could be to include variables such as sea surface temperature, rainfall, cloud cover, slope, aspect, reflectivity, wind speed and humidity in an eco-evolutionary model such as ours to make more accurate predictions for the fate of green sea turtle populations.

Another species with temperature-dependent sex determination that is at risk because of climate change is the tuatara (*Sphenodon punctatus*), a native New Zealand reptile that has also a relatively long generation time. Unlike green sea turtles, tuatara currently have small current population sizes and low levels of genetic variation. Tuatara have the opposite pattern of TSD, with female hatchlings at cold temperatures and male hatchlings at warm temperatures. A biophysical microclimate model predicts that under maximum warming forecasts for the 2080, almost all nests will produce 100% male hatchlings [57]. A shortage of males has even more immediate demographic consequences than a shortage of females. This is because warmer temperatures lead to a rise in females. Initially, this rise in females causes a rise in population size, since few males can mate with many females [58]. However, when temperatures continue rising and males become scarce, matings become scarcer as well, eventually driving the population to extinction. On the other hand, for cool temperatures, an increase in males leads to a steep population decline. In a study comparing the extinction risk for species with environmental sex determination under climate change based on a set of criteria, the tuatara received the highest risk score [2]. Because of their MF sex-determination system and the higher population size and levels of genetic variation, green sea turtles, which were not included in that study, would likely receive a somewhat lower risk score. However, our results show that even if we optimistically assume substantial heritable variation in relevant traits, there are limits to ability of this species to adapt to a warming climate.

## 5. Conclusions

To conclude, modelling green sea turtle population ecology and evolution shows some possible ways to support green sea turtle survival. The turtles may be able to adapt to climate change, if the CO_2_ emissions stay within a low to moderate range. However, model assumptions on heritability of traits and variance within the population remain to be empirically tested to ensure model accuracy. In the meantime, anthropogenic conservation measures may support population survival without compromising evolution.

## Author Contributions

Conceptualization, J.B. and C.B.; methodology, J.B.; software, J.B. and C.B.; validation, J.B., M.J.W. and C.B.; formal analysis, J.B and C.B..; investigation, J.B. and C.B.; resources, J.B.; data curation, J.B. and C.B.; writing—original draft preparation, J.B..; writing—review and editing, C.B. and M.J.W..; visualization, C.B..; supervision, C.B. and M.J.W.. All authors have read and agreed to the published version of the manuscript.

## Acknowledgments

We would like to thank Klaus Reinhold for comments on an earlier version of the manuscript. We thank Carola Best, Sarah Connors, and Kirsten Zickfeld for their help in providing and understanding temperature prediction data. Moreover, we would like to thank Michael Jensen for advice on green sea turtle biology, and finally the Theoretical Biology group for discussion.

## Appendix A Calculation of the TSD equation (equation 1)

*a* is the upper limit, and *b* is the constant of proportionality. They were calculated using the following data by Limpus [13]: at 29.3°C a 0.5 sex ratio of females is produced, so *t*_*piv*_ = 29.3. For temperatures going towards infinity, the sex ratio of females will go towards 1, which requires *a* = 1. At 28°C the sex ratio is 0.05 female [13], i.e.

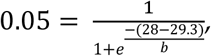

leading to

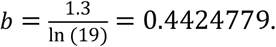

## Appendix B

**Figure A 1:**
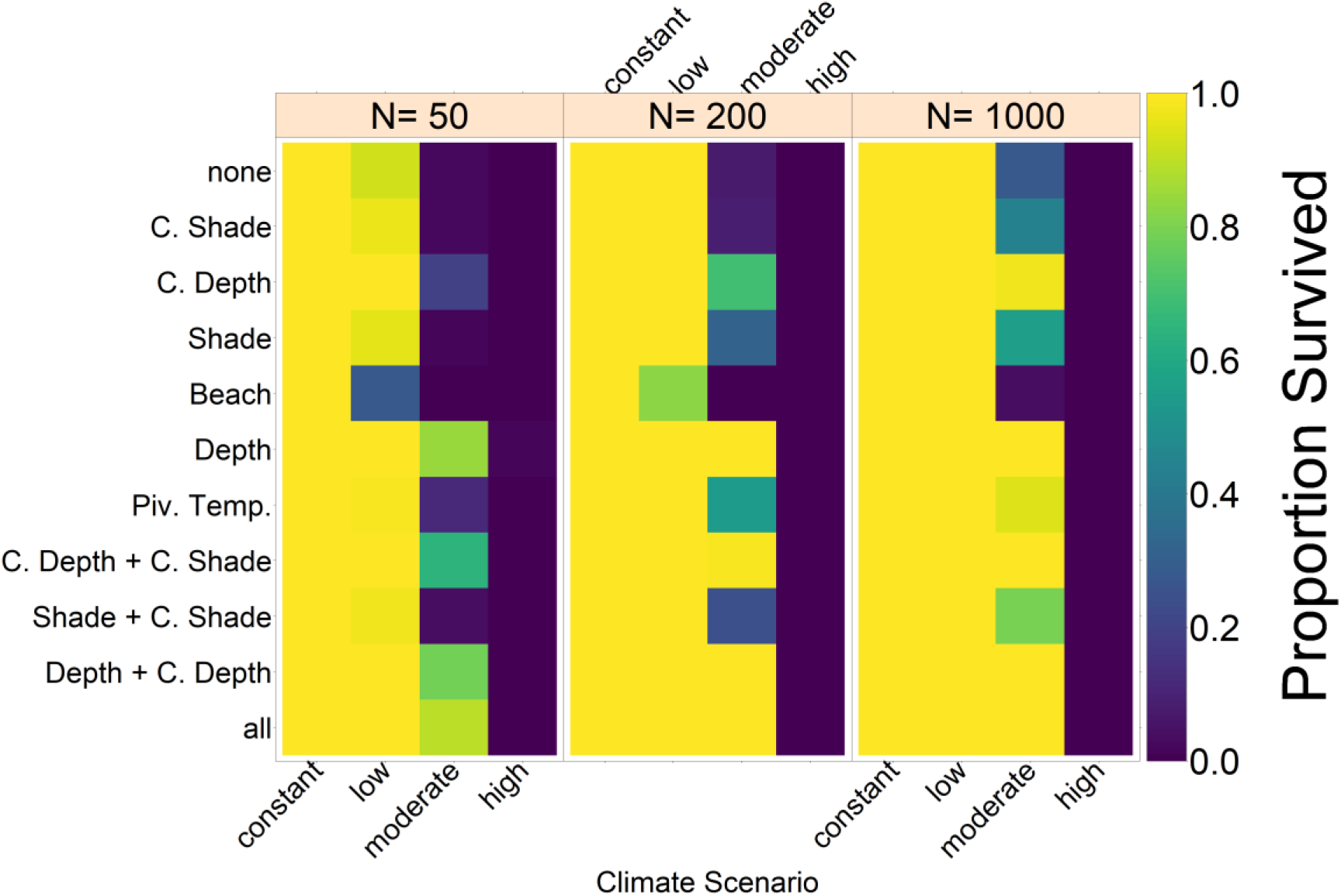
Effect of carrying capacity and thereby initial population size. Colors indicate the proportion of populations surviving to the year 2500. Note that the number of mating areas *g* was always half the carrying capacity to carrying capacity for this simulation. Varying the carrying capacity does not have a noticeable effect on population survival in the various setups.

**Figure A 2:**
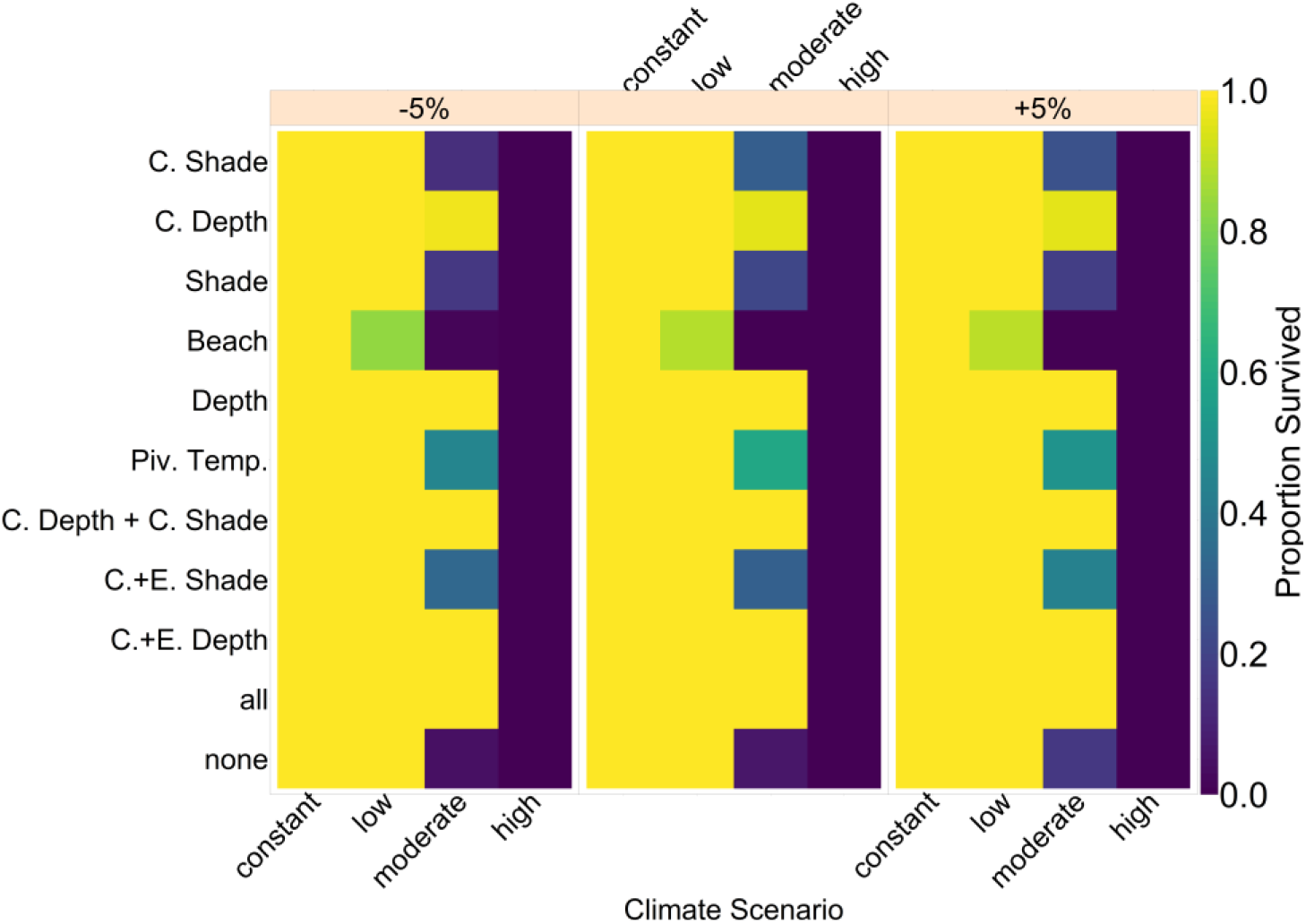
Robustness analysis for age at maturity. Results for varying parameters by ±5%. The y-axis shows each mechanism and any combinations thereof; the x-axis shows the parameters for each of the four climate scenarios. Colors indicate the proportion of replicates in which the population

**Figure A 3:**
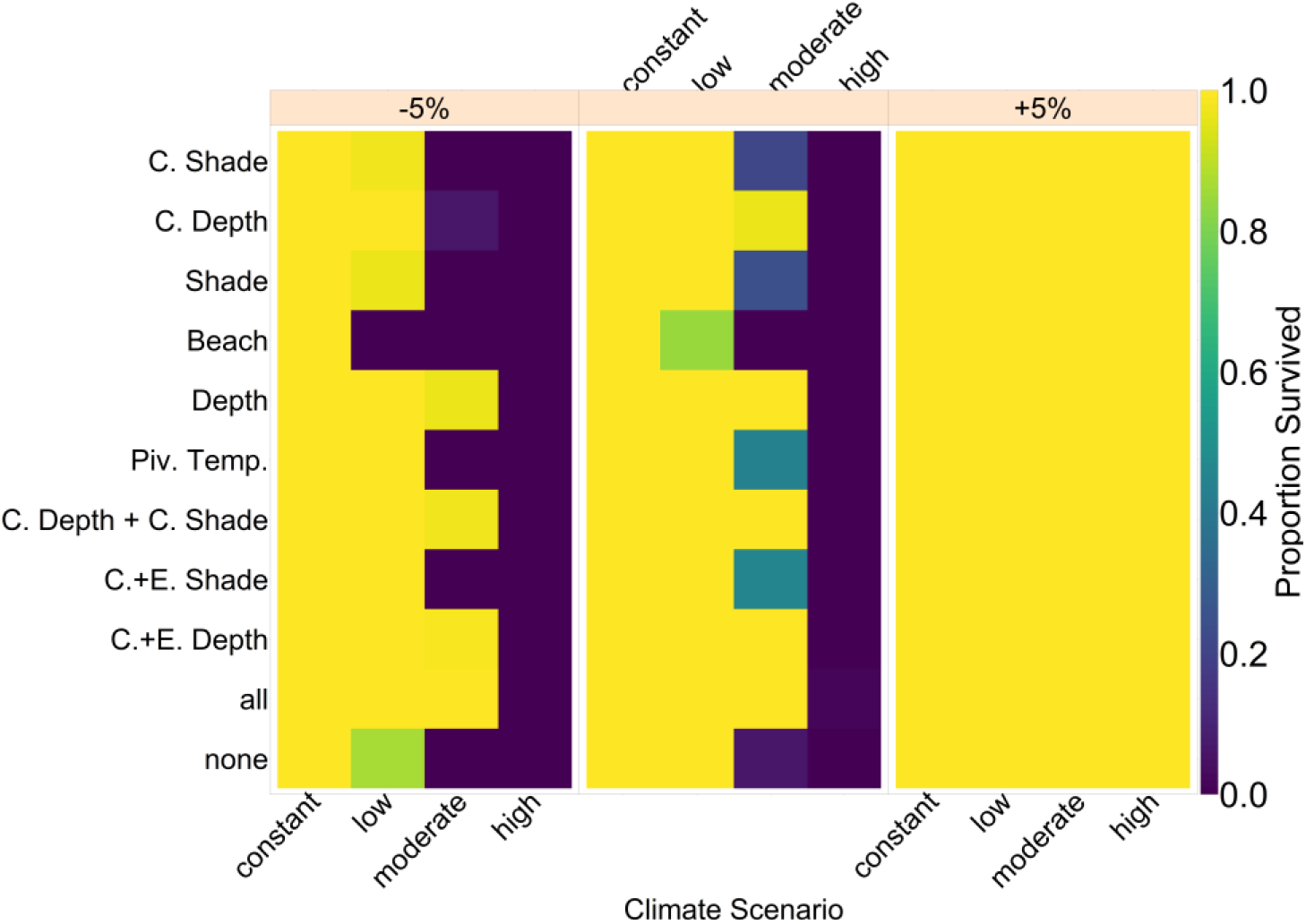
Robustness analysis for both juvenile and adult survival rates. Results for varying both parameters jointly by ±5%. The y-axis shows each mechanism and any combinations thereof; the x-axis shows the parameters for each of the four climate scenarios. Colors indicate the proportion of replicates in which the population survived up until year 2500.

**Figure A 4:**
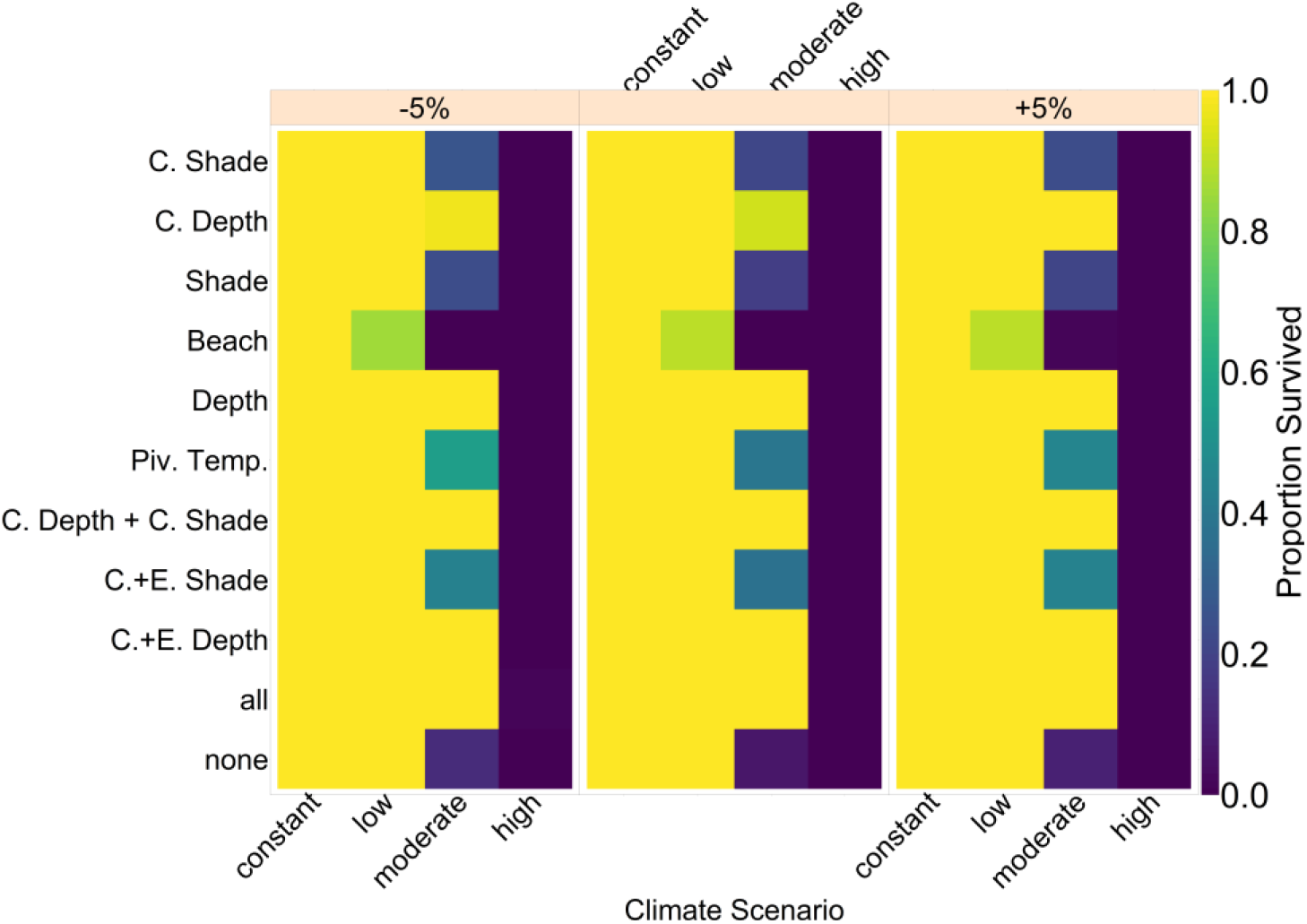
Robustness analysis for standard deviation of the Beta distibution representing mutation. Results for varying parameters by ±5%. The y-axis shows each mechanism and any combinations thereof; the x-axis shows the parameters for each of the four climate scenarios. Colors indicate the proportion of replicates in which the population survived up until year 2500.

**Figure A 5:**
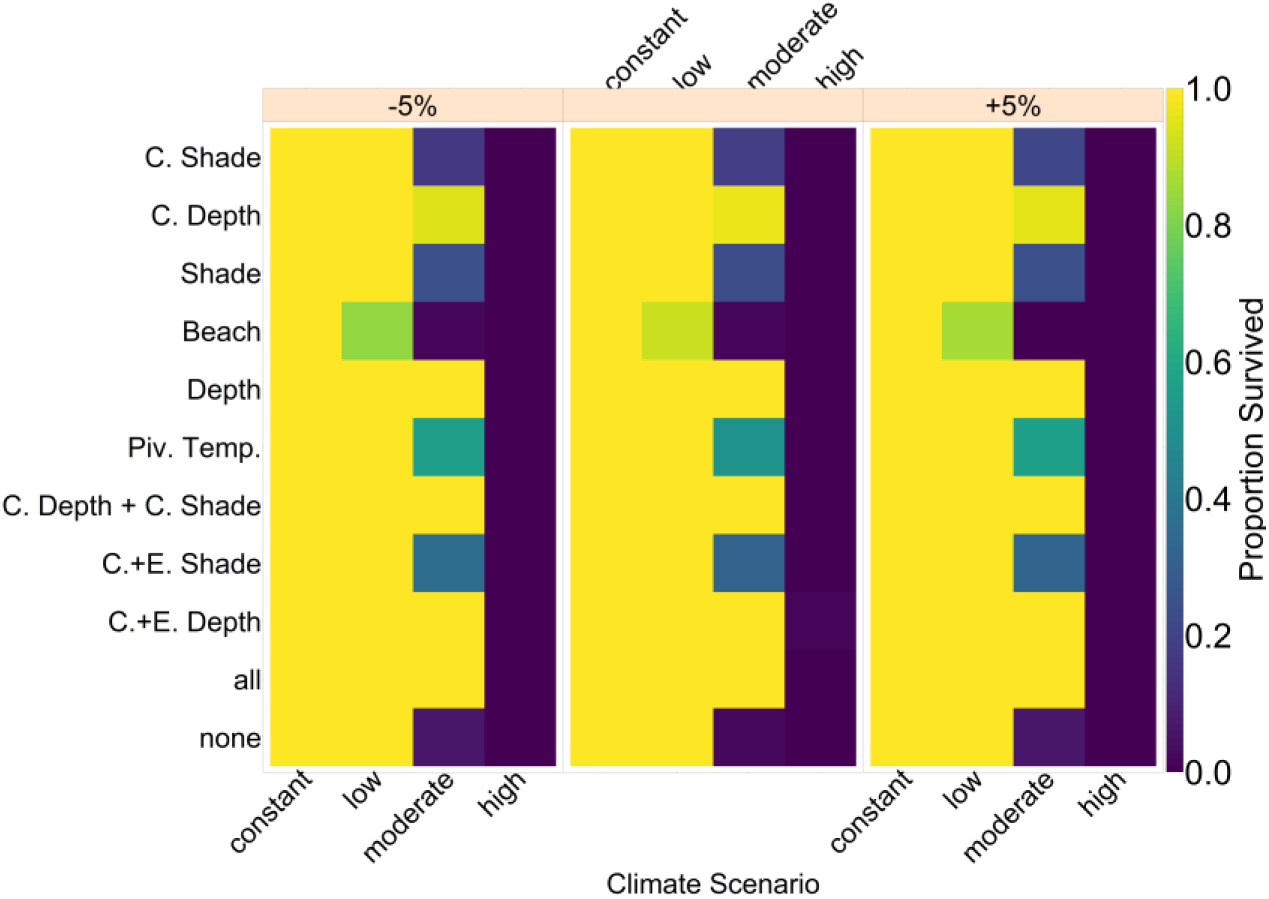
Robustness analysis for pre-industrial baseline nest temperature. Results for varying parameters by ±5%. The y-axis shows each mechanism and any combinations thereof; the x-axis shows the parameters for each of the four climate scenarios. Colors indicate the proportion of replicates in which the population survived up until year 2500.

**Figure A 6:**
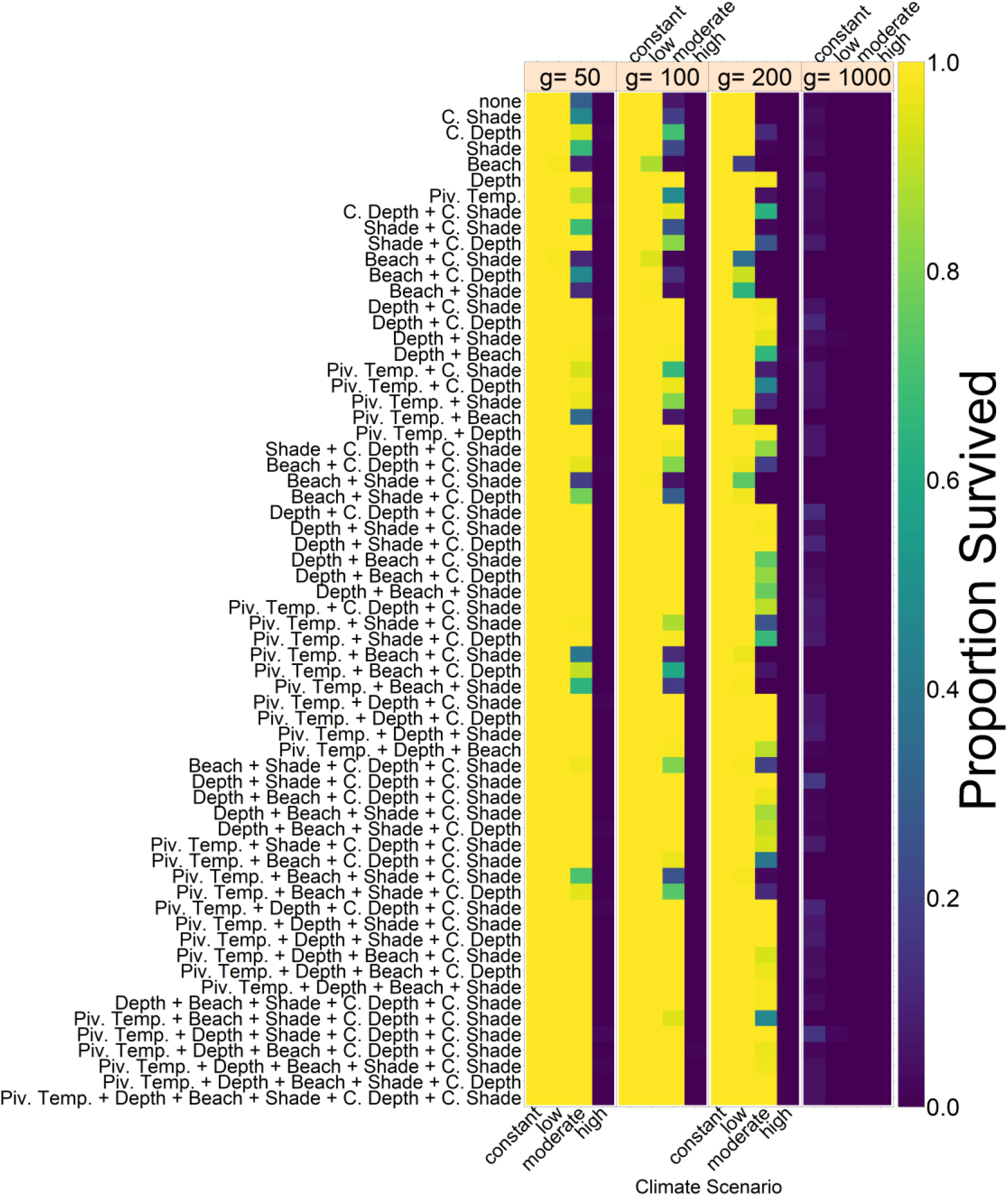
Effect of number of mating areas, *g*. The default value used for all other results is *g=100*. This Figure shows a summary for all possible combinations of trait evolution, philopatry and conservation measures. Colors indicate the proportion of populations surviving to the year 2500. The more mating areas there are, the less likely the population is to survive, since the probability of finding a mate decreases with increasing number of mating areas.

**Figure A 7:**
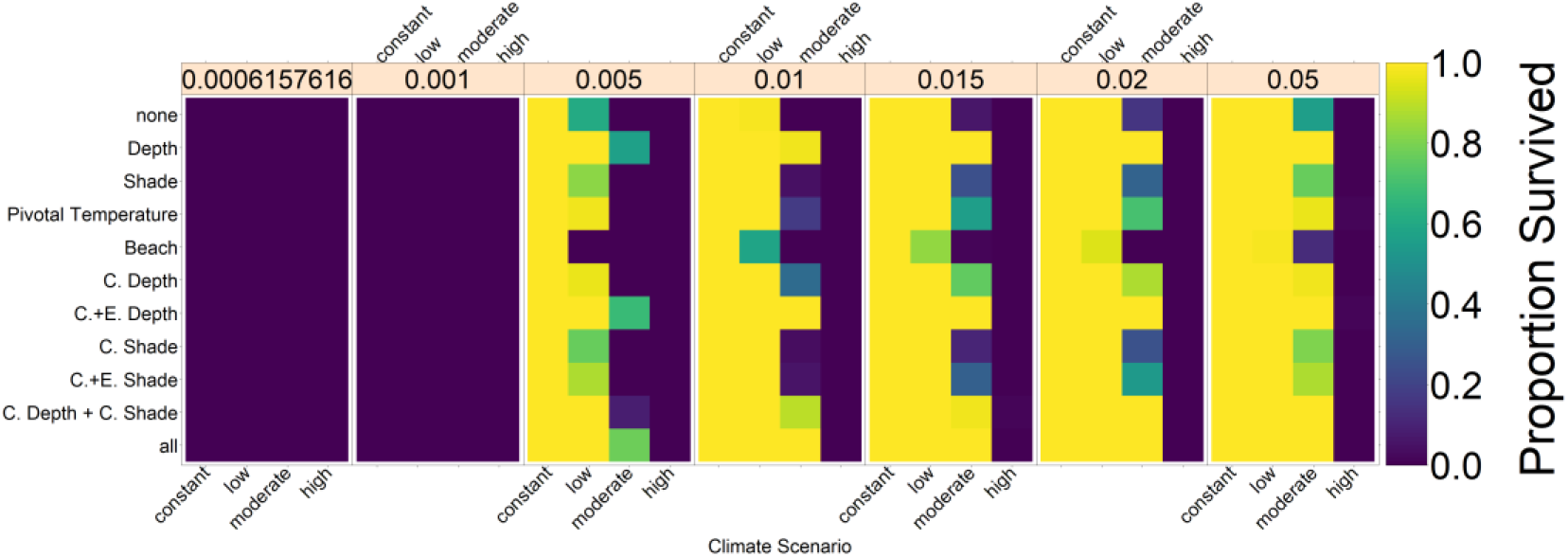
Robustness analysis for juvenile mortality, i.e. the survival probability from egg to mature adult. Colors indicate the proportion of replicates in which the population survived up until year 2500. The leftmost box shows survival rate based on Chaloupka [38], where juvenile survival probability was chosen to obtain stable populations, but with a different set of parameters and assumptions.

**Figure A 8:**
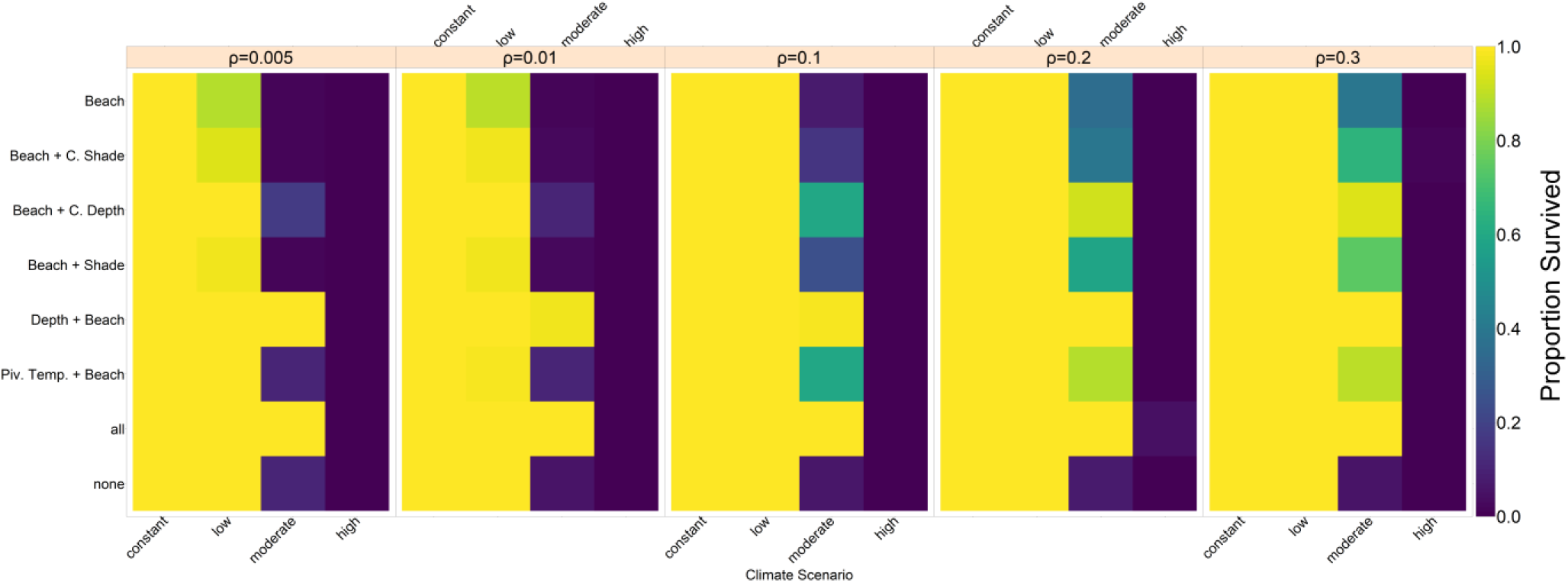
Effect of level of nest site philopatry 1-ϱ. Numbers in orange boxes indicate the ratio of females that do not return to their native beach. Colors indicate the proportion of populations surviving to the year 2500.

**Figure A 9:**
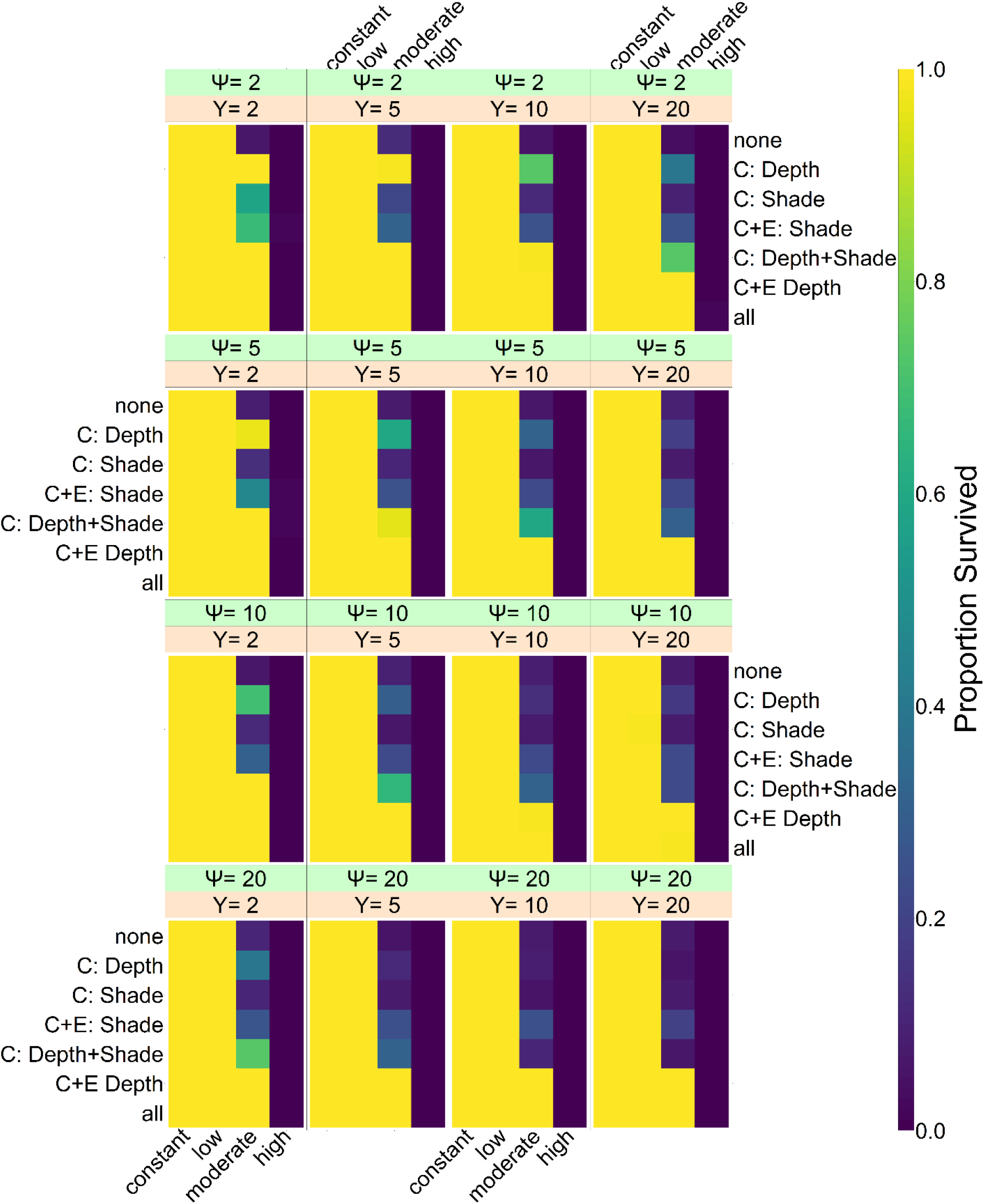
Effect of conservation measures. ϒ is the interval at which efforts are carried out, e.g. ϒ=10 means that the efforts are carried out every 10 years. Ψ is the frequency at which nests are manipulated, e.g. Ψ=2 means every 2^nd^ nest is manipulated. Colors indicate the proportion of populations surviving to the year 2500.

**Figure A 10:**
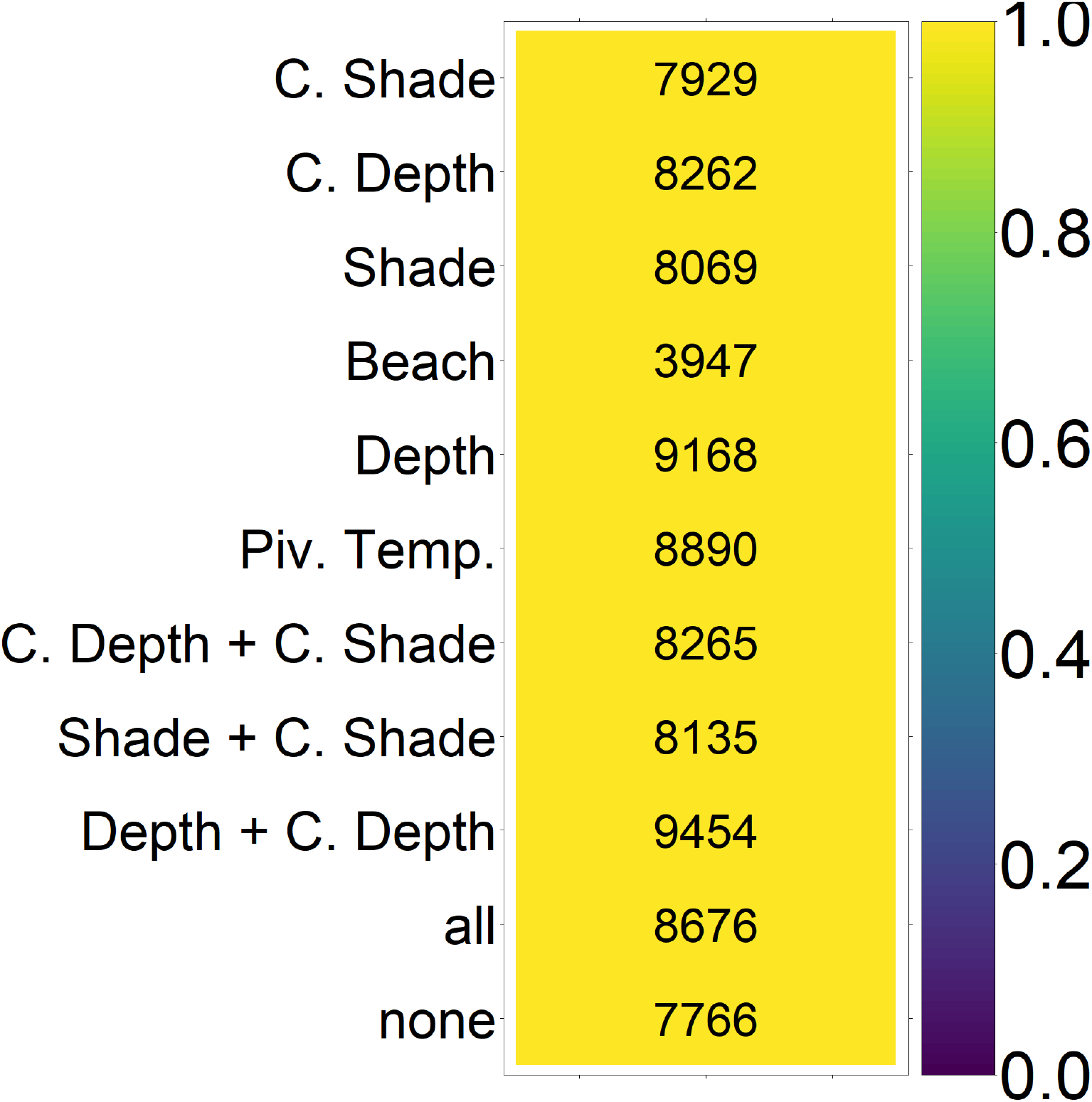
Results from the generation of a population for the southern Great Barrier Reef. Adult males were used as a source of migration to the nGBR for corresponding simulations. Colors indicate the proportion of replicates that survived for 1000 years. Numbers are the number of males retrieved for each setup.

